# Molecular insights from conformational ensembles via machine learning

**DOI:** 10.1101/695254

**Authors:** O. Fleetwood, M.A. Kasimova, A.M. Westerlund, L. Delemotte

## Abstract

Biomolecular simulations are intrinsically high dimensional and generate noisy datasets of ever increasing size. Extracting important features in the data is crucial for understanding the biophysical properties of molecular processes, but remains a big challenge. Machine learning (ML) provides powerful dimensionality reduction tools. However, such methods are often criticized to resemble black boxes with limited human-interpretable insight.

We use methods from supervised and unsupervised ML to efficiently create interpretable maps of important features from molecular simulations. We benchmark the performance of several methods including neural networks, random forests and principal component analysis, using a toy model with properties reminiscent of macromolecular behavior. We then analyze three diverse biological processes: conformational changes within the soluble protein calmodulin, ligand binding to a G protein-coupled receptor and activation of an ion channel voltage-sensor domain, unravelling features critical for signal transduction, ligand binding and voltage sensing. This work demonstrates the usefulness of ML in understanding biomolecular states and demystifying complex simulations.

**STATEMENT OF SIGNIFICANCE:** Understanding how biomolecules function requires resolving the ensemble of structures they visit. Molecular dynamics simulations compute these ensembles and generate large amounts of data that can be noisy and need to be condensed for human interpretation. Machine learning methods are designed to process large amounts of data, but are often criticized for their black-box nature and have historically been modestly used in the analysis of biomolecular systems. We demonstrate how machine learning tools can provide an interpretable overview of important features in a simulation dataset. We develop a protocol to quickly perform data-driven analysis of molecular simulations. This protocol is applied to identify the molecular basis of ligand binding to a receptor and of voltage sensitivity of an ion channel.

## INTRODUCTION

Molecular dynamics simulations of biological systems provide a unique atomistic insight into many important biological processes such as a protein’s conformational change between functional states, the folding of a soluble protein or the effect of ligand binding to a receptor. These systems can be extremely high dimensional with pairwise interactions between tens to hundreds of thousands of atoms at every snapshot in time. As system sizes, as well as the reachable timescales of simulations, have increased significantly over the last decade, a conventional simulation can now generate terabytes of raw data that needs to be condensed for human interpretation. Data-driven methods reduce the risk for researchers to overlook important properties in a simulation, or even misinterpret the computational experiment and introduce human bias in the analysis.

A system of N particles has 3N spatial degrees of freedom. Fortunately, restraints in the system due to, for example, the force field and steric hindrance, restrict biomolecules to only adopt a subset of all possible atomic rearrangements. Thus, the system typically moves on a manifold of much lower dimensionality than its actual number of degrees of freedom. Finding this manifold is not easy, and even when it is found it can still be difficult to *interpret* it. Researchers often seek *reaction coordinates* or *Collective Variables* (CVs) to describe important features of this manifold. An optimal set of CVs should help answering a specific question. For example, if the aim is to enhance sampling of transitions from one functional state to another, we typically seek the CVs which best describe the slowest motion of the system. In other situations we might seek answers to biophysically relevant questions by chasing subtle differences on the molecular level between slightly perturbed states. Examples include establishing the molecular signatures of a protein when bound to different ligands, considering different protonation states or the effect of a point mutation. Regardless of the biological problem at hand, analysis typically involves processing a large set of high dimensional data in search of important features. This type of dimensionality reduction problem may be addressed with Machine Learning (ML) methods.

The increasing amounts of data, as well as limited time and resources for processing and analysis, is not only a challenge in the field of biomolecular simulations. ML methods have gained enormous interest in recent years and are now applied in a wide range of research areas within biology, medicine and health care (1–3) such as genomics (4), network biology (5), drug discovery (6) and medical imaging (7, 8). In molecular simulations, such methods have for example eminently been used to enhance sampling by identifying CVs or the intrinsic dimensionality of biomolecular system in a data-driven manner (9–30), as an interpolation or exploratory tool for generating new protein conformations (23, 31, 32), as well as providing a framework for learning biomolecular states and kinetics (11, 33, 34). However, many tools borrowed from ML, most notably nonlinear models such as Neural Networks (NN), are criticized for their black-box resemblance which obstructs human interpretable insights (1–3, 5). Therefore, making this opaque black box transparent is an active area of research (35, 36).

In this study we have demonstrated how to learn ensemble properties from molecular simulations and provide easily interpretable metrics of important features with prominent ML methods of varying complexity, including principal component analysis (PCA), random forests (RFs) and three types of neural networks: autoencoders (AEs), restricted Boltzmann machines (RBM) and multilayer perceptrons (MLP). For different types of datasets, we considered supervised methods, where each simulation frame belongs to a known class, as well as unsupervised methods, where information is extracted from unlabelled data. In order to show how the methods perform under various circumstances, we first evaluated them on a toy model designed to mimic real macromolecular behavior, where functional states were defined by state-dependent random displacement of atoms. This approach enabled us to understand and illustrate the different methods’ properties and shortcomings in a quantitative manner. In particular, we find that all methods perform equally well on the toy model’s simple setup, in which optimal inverse interatomic distances are used as an input. However, **while supervised methods are still able to identify important features for a more complex input such as Cartesian coordinates, unsupervised methods fail to do so**.

We further applied our protocol for identifying key features of three biologically interesting processes that involve systems sampled using extensive conventional MD or enhanced sampling techniques: the conformational rearrangements occurring within the C-terminal domain of the Ca^2+^-bound soluble protein calmodulin, the effect of ligand binding to a G protein-coupled receptor (GPCR) and the response of an ion channel voltage-sensor domain to a change in the transmembrane potential. In all three cases, the ML methods pinpointed molecular features that are known to be important for the function of these proteins. In general, our results demonstrate the ability of ML methods to reveal valuable insights into biomolecular systems. We anticipate that this straightforward, yet powerful approach, can become useful for many researchers when demystifying complex simulations.

## METHODS

### Principal component analysis

**Principal Component Analysis (PCA) converts a set of input features,** *X*, *to a set of orthogonal linearly uncorrelated variables*, *T*, **corresponding to the eigenvectors of of** *X^T^X* **with normalized eigenvalues** *λ* **(37). The eigenvectors with the highest eigenvalue covers the largest variance possible in the dataset and is called the first principal component (PC), the second PC the largest variance in the orthogonal space of the first component, etc … The feature importance**, *R*, **was taken to be the projection of the first PCs and eigenvalues onto the original input features, up to a specified threshold in accumulated variance (Fig. 1 A):** 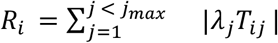, **where** *i* **is the index of the input features and** *j* **is the index of the PCs**. *j_max_* **was determined by three different approaches: 1) with a fixed number of PCs, 2) so that the** 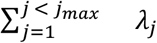 **was below a certain threshold or 3) set to the first** *λ_j_* **which fulfilled the condition** *λ_j-_/λ_j_* ≥10. Finally, the importance was normalized between 0 and 1.

**Figure 1.**
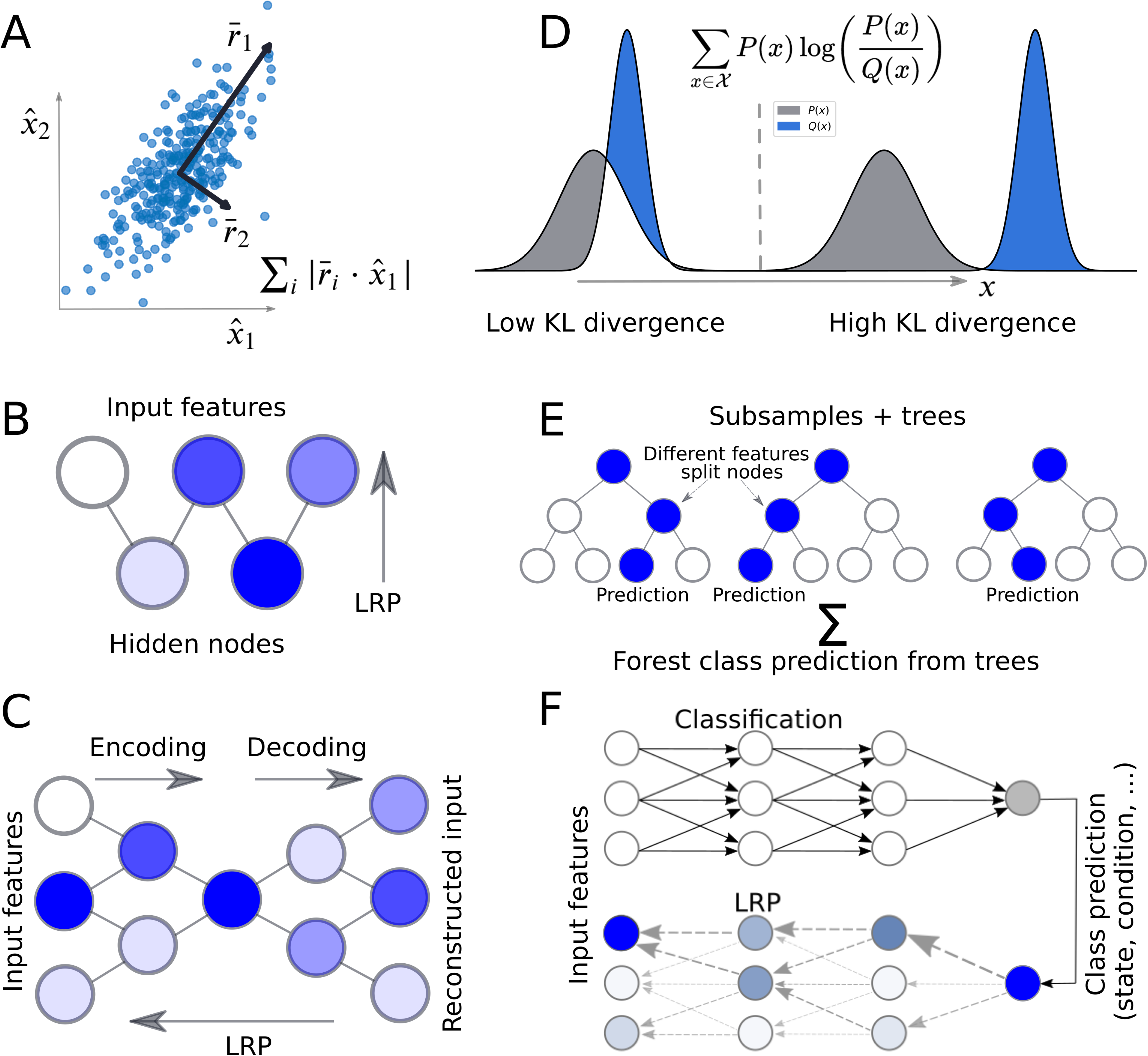
Machine learning models used in this study. (A) Principal Component Analysis (PCA) converts the input features to an orthogonal set of linearly uncorrelated variables called principal components (PCs) through an orthogonal transformation. The first component covers the largest variance possible. The feature importance is taken to be the components’ coefficients of a feature times the variance covered by (i.e. the eigenvalues) the components. (B) Restricted Boltzmann Machine (RBM): a generative stochastic neural network trained to maximize the likelihood of the data using a graphical model with a layer of hidden nodes connected to the input nodes. Important features can be derived using Layer Relevance Propagation (LRP), an algorithm originally developed for image classification problems (39). (C) Autoencoder (AE): a generative neural network trained to reconstruct the features through a set of hidden layers of lower dimensionality. Important features can be derived using LRP. (D) Kullback-Leibler (KL) divergence (also called relative entropy): this metric computes the difference between two distributions *P*(*x*) and *Q*(*x*) along every individual feature. We compute the KL divergence between one class and all other classes as an indication of the importance of a specific feature. (E) Random Forest (RF) classifier. A prediction is taken as an ensemble average over many decision trees. Average relevance per feature is computed as the mean decrease impurity (44). (F) A Multilayer Perceptron (MLP) is a feedforward artificial neural network with fully connected layers. Important features can be derived using LRP.

### Restricted Boltzmann machine

A restricted Boltzmann machine (RBM) (38) is a generative stochastic neural network trained to maximize the likelihood of the data using a graphical model with a layer of hidden nodes connected to the input nodes (Fig. 1 B). We used an implementation from scikit-learn of a Bernoulli RBM. Training was performed with stochastic maximum likelihood. To find important features, **Layer-wise relevance propagation (LRP; explained in the following section)** was performed from the output of the hidden layer to the input layer.

Note that in all computations, the input features were scaled between values of 0 and 1. The RBM used in this study assumes a probabilistic interpretation of the input features with values between 0 and 1, where the upper limit indicates that a specific input node is active. When feeding inverse distances to the RBM, larger input values correspond to residues in contact with each other, this model makes physical sense as long as the values are scaled properly. Cartesian coordinates, even after scaling, do not fulfill the assumptions made about the input features and have been taken into consideration only for the sake of consistency.

### Layer-wise relevance propagation (deep Taylor decomposition)

**Layer-wise relevance propagation (LRP),** or deep Taylor decomposition when applied to networks with ReLU activation (39), is a method originally developed for image classification problems with the aim to produce a decomposition (an explanation) of a prediction that can be visualized in the same way as the input data. Given a trained neural network, it is possible to perform a backward pass through the network (Fig. 1 F), **keeping track of which nodes contributed to a certain classification decision. More specifically, for a prediction with input features** *X*, **true labels** *T* **and a network output** *T*′, **the relevance vector**, *R*, **is initialized as** *R_i_*, = *T_i_T’_i_*. **The relevance is iteratively propagated from nodes in one layer (index** *k***) to the nodes in a previous layer (index** *j***) with the following iterative rule:** 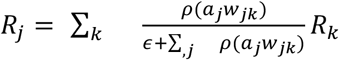. **Here,** *a_j_*, **is the activation of the current node,** *ϵ* **is a small increment set to** 10^-9^, *w_jk_* **are the weights connecting the two layers and** *ρ* **is a function which transforms the weights. For layers of bounded values, such as the input layer if** *X* **is scaled between fixed values**, *ρ = a_j_W_jk_* ‒ *l_j_max*(0, *w_jk_*) – *h_j_min*(0, *w_jk_*) **where** *l_j_* **and** *h_j_* **are the layer’s lower and upper bounds. For unbounded values**, *ρ* = *a_j_max*(0, *w_jk_*). The importance of a feature was derived by computing the average relevance over all input samples in the training set and normalizing the importance for all features and frames to have an upper bound of 1. Unlike other methods in this study, it is possible to compute the importance of a feature for a specific simulation frame in this way.

### Autoencoder

An autoencoder (AE) is a generative neural network trained to reconstruct the input through a set of hidden layers of lower dimensionality (Fig. 1 C) (40). In short, the model can be broken down into two parts: encoding and decoding. In the encoding part, the layers decrease in size causing the network to ignore noise during training. In the decoding part the layers increase in size up to the output layer, which consists of reconstructed input features.

The AE was implemented by training a scikit-learn MLP-regressor (41) with the same data as input and output. The encoding layers and the decoding layers were always set to be of the same shape. To find important features, LRP was performed from the reconstructed output to the actual input.

### Kullback Leibler divergence

The Kullback Leibler (KL) divergence (Fig. 1 D) (42), also called relative entropy, is a measure of the *p*(*x*) difference between two distributions *P*(*x*) and *Q*(*x*) along a feature X: 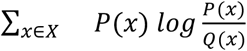. The importance of a feature was set to be a symmetric version of the KL divergence, 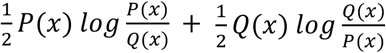, between one state and all other states along that feature. All importance scores were normalized between 0 and 1.

### Random Forest classifier

A random forest (RF) classifier (43) (Fig. 1 E) is an ML model in which a prediction is taken as an ensemble average over many decision trees, **each tree fitted to a subsample of the dataset**. Important features were identified by computing the forest’s normalized mean decrease impurity (44), using scikit-learn’s (41) implementation. **Mean decrease impurity is essentially a measure of how often a feature is used to split nodes in the underlying decision trees. It is computed by identifying all nodes split by that feature and averaging the metric** *ρΔi* **over all trees in the forest, where** *ρ* **is the fraction of samples reaching the node and** *Δi* **is the decrease in impurity by the split (45). In this study we used the Gini impurity, computed for a dataset with** *N* **classes as** 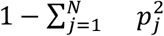, **where** *p_j_* **is the probability of a sample in class *j*reaching that node.** To compute the feature importance for a specific state, the RF was alternatively used as a one-vs-the-rest binary classifier trained to distinguish one state from all the others.

### Multilayer perceptron classifier

A Multilayer perceptron (MLP) classifier (Fig. 1 F) is a feedforward artificial neural network with fully connected layers (42). An implementation from scikit-learn (41) of an MLP classifier was used throughout this study, trained with the Adam solver (46). To find important features, LRP was performed from the output to the input layer.

### Toy Model

To construct model systems, atoms were randomly placed in a box to set up an initial configuration. To define states, a set of atoms unique to every state, typically few atoms compared to the size of the system, were labelled as important and displaced relative to the initial configuration. The toy model equivalent of simulation frames in a trajectory were generated by displacing every labeled atom’s position considering uniform noise and randomly rotating the entire system around the origin. In silico systems are typically rotation invariant but aligning different configurations is often not trivial. In addition to the number of atoms and frames in the generated trajectory, the number of states, the number of important atoms per state and the strength of the noise were all configurable parameters of the toy model.

Moreover, we constructed two kinds of displacements: a linear displacement of atoms from their original position (Fig. 2 A) as well as a nonlinear displacement method (Fig. S1 and SI Methods). In the linear method, the magnitude and direction of the displacement was the same for all atoms. In the nonlinear method, one atom per state was displaced from its equilibrium position, the size and the direction of the displacement depending on the state index. Subsequent atoms were rotated around the position of the previous atom in the same state (see SI Methods). From a biomolecular perspective, this can be seen as a model of how interresidue contacts are broken in a linear (for example with the displacement of one subdomain away from another) or nonlinear manner (for example with the twist of a helix).

**Figure 2.**
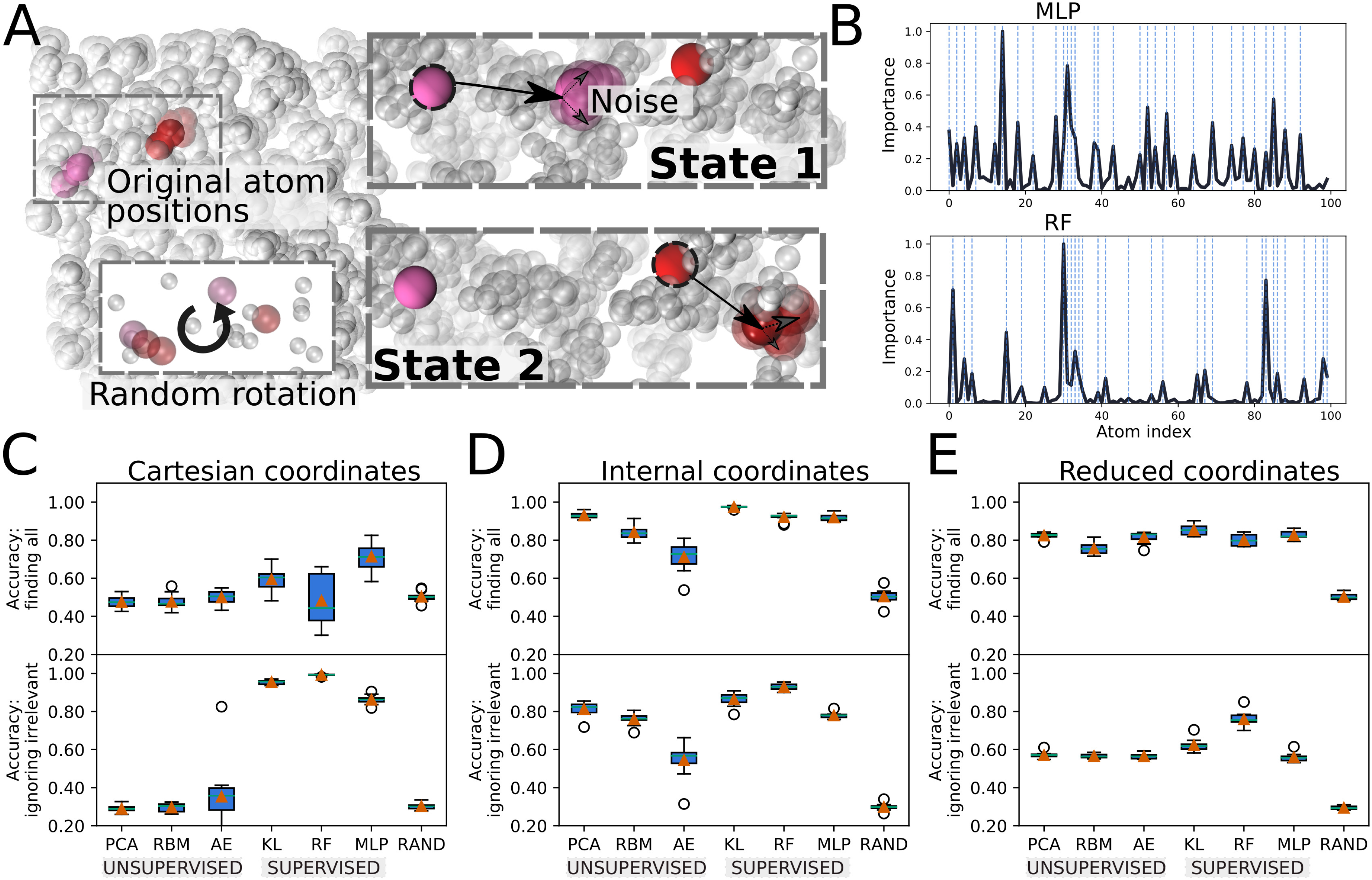
(A) Toy model used to benchmark different ML approaches to extract important features from simulation data. Atomic coordinates are randomly generated and a subset of the atoms, unique to every state, is displaced linearly from their initial positions. Artificial simulation frames are generated by adding noise to all atoms’ positions. Since only the relative and not the absolute positions of atoms are of significance in real biological systems, the system is rotated randomly around the origin. (B) Importance per atom for single instances of the toy model. The index of the displaced atoms are highlighted as dashed vertical lines, coinciding with all peaks in importance in the case of a Multilayer Perceptron (MLP), and some of them for a Random Forest (RF). (C-E) Boxplots of the performance of the different methods using either Cartesian coordinates (C), the full set of inverse interatomic distances (D) or a reduced set of inverse interatomic distances (E) as input features sampled over different instances of the toy model with linear displacement and 10% noise level. A high accuracy at finding all atoms signifies that every displaced atom has been identified as important and that other atoms have low importance. A high accuracy at ignoring irrelevant atoms signifies that only displaced atoms, although not necessarily all of them, have been marked important (Methods and Fig. S10). The best performing set of hyperparameters found after benchmarking every method (Fig. S4-S9) have been used. RAND stands for random guessing.

The magnitude of the displacement was 0.1 and the strength of the noise was 0.01 except for Fig. S2 in the Supporting Material, where it was increased to 0.05. 1200 frames were generated for each state. In order to obtain statistics, we generated 10 toy models with 100 atoms in a box with a side length stretching from −1 to 1, **with the exception of Fig. S3 where we evaluated systems of varying size**. In total there were 3 states with 10 displaced atoms per state. For every ML method we computed the performance for different combinations of hyperparameters (Fig. S4-S9). For every instance of the toy model, the feature importance was computed 10 times. The parameters which gave the best performance were included in Fig. 2, S1 and S2.

When using inverse distances as features, the relevance per atom was computed by summing the average importance of all distances involving an atom and normalizing the values between 0 and 1. To take into account the smallest number of input features possible without losing information regarding the positions of the atoms, we also constructed a reduced set of internal coordinates. We did so by including the distances from atom i to atoms i+1, i+2, i+3 and i+4 only; this is akin to triangulation and suffices to reconstruct the coordinates of the entire system. Cartesian coordinates generated 3N input features. To compute the importance per atom, the importance of every triplet of xyz coordinates was summed and normalized.

### Accuracy quantification

We used two scores to estimate a method’s performance on the toy model. Given a measured distribution of importance 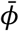, and a true distribution, 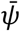, we computed the accuracy at *finding all* important residues as: 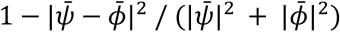. This mean-square-error (mse) based metric gave high scores to methods which identified all displaced atoms in the toy model and could tolerate some noise. The *ignoring irrelevant* accuracy score, designed to give high scores to methods which only identified truly important atoms without necessarily identifying all of them, was defined as 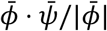. To illustrate the scoring method, the accuracy scores of various distributions are highlighted in Fig. S10.

### Calmodulin (CaM) analysis

The dataset consisted of six CaM C-terminal states extracted with spectral clustering from a mix of regular MD (3600 ns) and temperature enhanced simulations (570 ns replica exchange MD and 460 ns replica exchange solute tempering) (47). We used inverse Cα distances between C-terminal domain residues and a total of 7600 frames, yielding a dataset size of 7600 x 2145. The data was first shuffled in blocks of 100 frames and filtered prior to feature extraction. Only residue pairs with Cα distance less than 1.0 nm in at least one frame and more than 1.0 nm in another were kept. This cutoff-filtering decreased the number of features to 688. All features were normalized with scikit-learn’s (41) min-max scaler. To pick the number of components in PCA, a variance cutoff of 75 % was used. The AE and MLP both had hidden layers of 120 and 100 nodes, respectively. Both were trained with an Adam optimizer and used RelU activation functions. The RF classifier used 500 estimators and was trained with a one-vs-rest approach to obtain important features for every individual state. We used 3-fold cross-validation and averaged the residue profile over 5 independent iterations.

### G protein-coupled receptor (GPCR) analysis

The dataset contained the output coordinates from every tenth short 10 ps trajectory initiated from points along the most probable transition path between β2AR’s active and inactive state. Trajectories from every tenth of the 200-300 iterations of a string of swarms simulation (10) were used. Inverse Cα distances between all 284 protein residues were normalized with scikit-learn’s min-max scaler and were chosen as input features.

All PCs until the corresponding eigenvalue differed by at least an order of magnitude compared to the next eigenvalue were used to perform PCA. An RF classifier with 1000 estimators was trained to discriminate between ligand bound and unbound frames. In total 2674 frames were used in training with 4-fold cross-validation. The final importance profiles were averaged over 30 independent iterations.

### Voltage sensor domain (VSD) analysis

**Snapshots corresponding to configurations making up five metastable states along VSD activation were extracted from a metadynamics trajectory taken from our previous study (48, 49).** The five states were previously identified using the network of salt bridges between the S4 positive residues and their negative counterparts as a collective variable (50). The overall length of the trajectory was 12175 frames with 1087, 1900, 2281, 4359, and 2548 frames corresponding to E, *Δ, Γ*, B and A states, respectively. Inverse heavy-atoms distances between all VSD residues were used as input features (13041 in total). Distances larger than 0.7 nm or smaller than 0.5 nm throughout the entire trajectory were filtered out yielding a final number of 3077 input features. To identify important residues, an RF classifier was trained with 100 estimators and 3-fold cross-validation. The final residue importance profile was averaged over 5 independent iterations.

### Software availability

Software used in this study is available for download in Delemottelab’s GitHub repository (https://github.com/delemottelab/demystifying).

**Table 1.**
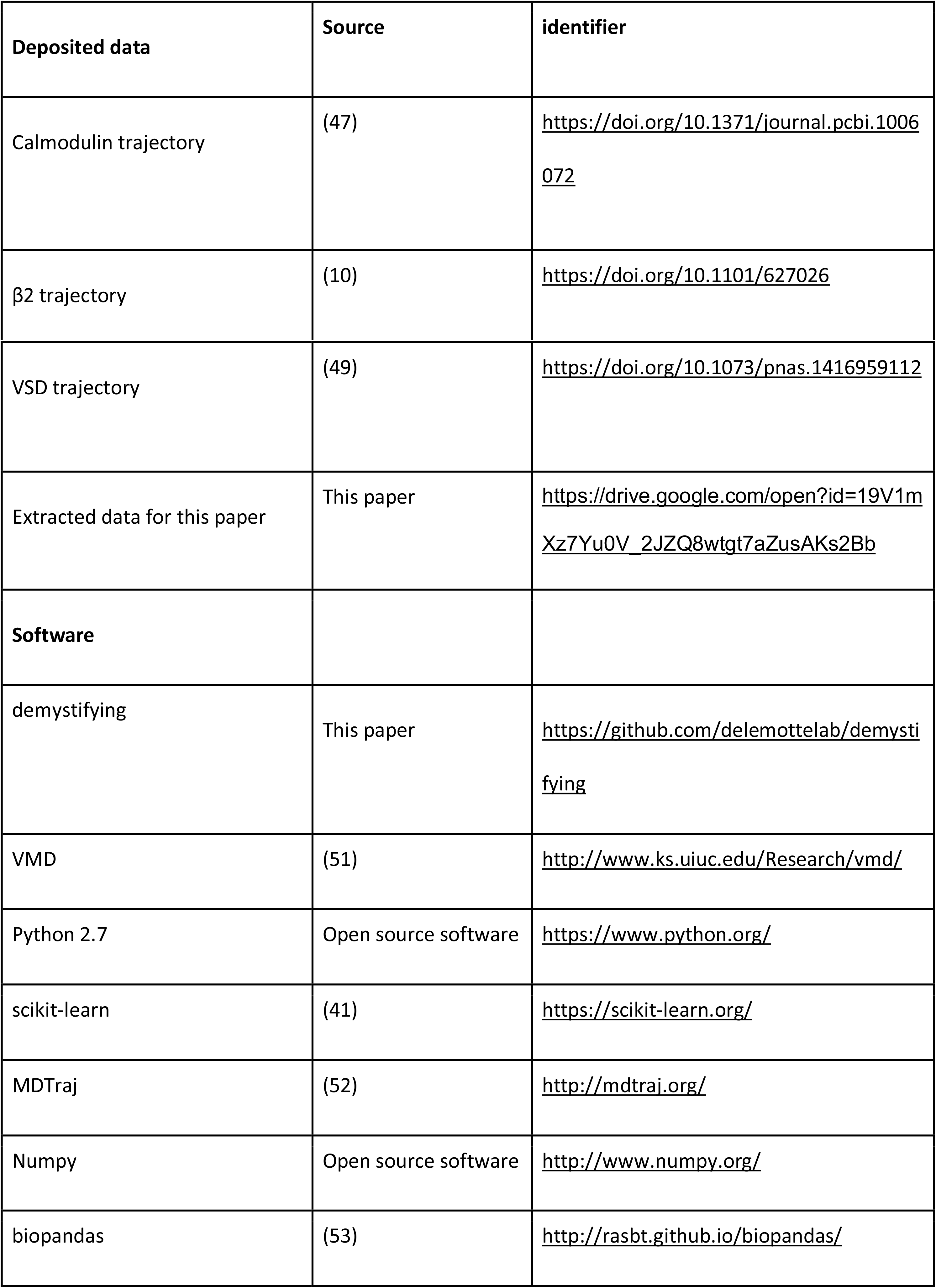
Resources - Data and Software

| **Deposited data** | **Source** | **identifier** |
| --- | --- | --- |
| Calmodulin trajectory | (47) | https://doi.org/10.1371/journal.pcbi.1006072 |
| β2 trajectory | (10) | https://doi.org/10.1101/627026 |
| VSD trajectory | (49) | https://doi.org/10.1073/pnas.1416959112 |
| Extracted data for this paper | This paper | https://drive.google.com/open?id=19V1mXz7Yu0V_2JZQ8wtgt7aZusAKs2Bb |
| **Software** |  |  |
| demystifying | This paper | https://github.com/delemottelab/demystifying |
| VMD | (51) | http://www.ks.uiuc.edu/Research/vmd/ |
| Python 2.7 | Open source software | https://www.python.org/ |
| scikit-learn | (41) | https://scikit-learn.org/ |
| MDTraj | (52) | http://mdtraj.org/ |
| Numpy | Open source software | http://www.numpy.org/ |
| biopandas | (53) | http://rasbt.github.io/biopandas/ |

## RESULTS AND DISCUSSION

### Benchmarking with a toy model provides quantitative measures of ML methods’ performance and guidance for analysis of real systems

Biomolecular systems consist of a large number of atoms or molecules. At each functional state, these fluctuate around their equilibrium positions due to thermal motion. As time progresses, a typical biomolecule transitions between a number of metastable *states*, either spontaneously, or because of changes in its environment. For real systems, identifying important features of such metastable states is desirable, and ML offers data-driven methods for this purpose.

To evaluate the performance of a certain method, one needs to truly know how the states are defined. We have designed and used a toy model that mimics a biomolecular system and for which we are able to control the parameters describing states (Fig. 2 A). In short, the atoms were first randomly placed within a box at their equilibrium positions. To define new states, certain atoms were displaced from their original positions (Fig. 2 A) in either a linear (Fig. 2 C, D) or nonlinear manner (Fig. S1, Methods and SI Methods S1). Artificial simulation frames were then generated by adding noise to the atomic coordinates. The accuracy of a method at *finding all* displaced (relevant) atoms was evaluated using a metric based on the mean-squared error between the estimated and the true feature importance profiles. In addition to this, we measured the ability of a method to *ignore irrelevant* atoms (i.e. minimizing false positives) (Methods). The way these scoring methods describe known distributions is shown in Fig. S10. In this way, the toy model allowed us to test and quantify how successfully a method identified true positives. We completed this analysis by evaluating the performance of the different methods when considering Cartesian or internal coordinates as input features, in a full or reduced set (Fig. 2 C-E) and at different noise levels (Fig. S2).

Given a trajectory consisting of many atoms and frames, with none or very little prior knowledge of the system, a widespread first approach in the biomolecular simulation community would be to apply principal component analysis (PCA) (Fig. 1 A) to all atomic coordinates and evaluate the ability of the first principal components (PCs) to identify the features that contributed most to the variance in the data. Fig. 2 C and S1 illustrate some major drawbacks of this unsupervised approach. Since PCA intrinsically performs a linear mapping of input features to a low dimensional representation (37), the performance of the method is highly dependent on the choice of input coordinates (13). For example, PCA failed to detect the important features of the toy system and even performed worse than random guessing when using raw, unaligned Cartesian coordinates as input (Fig. 2 C, S1 A). To remedy this, we performed PCA on internal coordinates (Fig. 2 D), namely inverse interatomic distances, either using the full set of all pairwise distances or a reduced set (Fig. 2 E) only connecting an atom to the four atoms before and after it in the sequence (Methods). The interatomic displacement that was performed to construct the states is easily described by such input coordinates, and PCA was able to successfully identify all the correct atoms for this simpler setup of the toy model (Fig. 2 D).

In addition to PCA, two alternative unsupervised learning methods were evaluated: a Bernoulli restricted Boltzmann Machine (RBM) (Fig. 1 B) (38) and an autoencoder (AE) (Fig. 1 C) (40). For every set of input features and labels, important features for a single classification decision were derived using layer-wise relevance propagation (LRP) (Fig. 1 F and Methods) (39). RBM and AE were both poor at distinguishing important features from irrelevant ones using Cartesian coordinates (Fig. 2 C). They were able, however, to perform with higher accuracy for linear as well as nonlinear toy model displacements using internal coordinates (Fig. 2 D and S1 B). While AE is suggested to be a promising unsupervised method for datasets that require nonlinear transformation, our results show that it **has a performance similar to or at best marginally better than PCA for Cartesian coordinates and a reduced set of internal coordinates, as well as for systems of smaller size (Fig. 2, S3). For the full set of internal coordinates from larger systems, PCA performs better (Fig. 2, S3).** This outcome indicates that AE is difficult to train on datasets of large dimensionality such as biomolecular simulations, and that to use this method successfully one should reduce the number of input features to a minimum.

The main limitation of PCA and other unsupervised learning techniques is that when the conformational states are known and each frame is labeled according to the state it belongs to, they fall short in utilizing this information. Supervised learning techniques, on the other hand, are designed for this purpose. In principle, supervised ML should be able to identify even the most subtle differences between states. We have applied three methods of different character and complexity (Fig. 2 C-E, S1 and S2) that take known class labels into account. As a simple first step, we computed the Kullback Leibler (KL) (Fig. 1 D) divergence (42) between one state and all other states for every individual input feature. A high KL divergence indicates that the feature is good at differentiating between states and hence important. Next, we trained a Random Forest (RF) (Fig. 1 E) classifier (43) to discriminate between states and computed feature importance using mean decrease impurity (44). The third method we evaluated was a Multi-layer Perceptron (MLP) classifier (Fig. 1 F), a prototypical neural network (42), in combination with LRP (39).

Computing the KL divergence between atomic coordinates does not transform the features in any way, and therefore failed to identify important atoms from the raw Cartesian coordinates (Fig. 2 C), especially when the noise in the toy model was strong (Fig. S2 A). On the other hand, it did not inaccurately score irrelevant atoms. RF also performed well for only one of the two accuracy metrics, specifically to ignore irrelevant atoms and map out some important ones, albeit not all of them (Fig. 2 B, C and S1 A). **This means that the classifier has successfully learned to predict states without taking all relevant features into account.** In many situations, this is an appealing characteristic; RF and KL provide quick overviews with a close-to-minimal set of features to describe the system and give highly interpretable importance profiles. This, however, comes at the risk of ignoring other significant features in the data. MLP, which has the ability to approximate nonlinear classification functions due to its multi-layer architecture and use of activation functions, successfully identified the majority of the important features from unaligned Cartesian coordinates (Fig. 2 B, C and S1 A). Its performance, compared to the other methods, was found to be superior when states were defined by nonlinear instead of linear displacement between atoms (Fig. S1 A) or after increasing the strength of the noise in the toy model (Fig. S2). On the other hand, it was not as good at ignoring irrelevant features, as it inaccurately scores some non-displaced atoms as relatively important by mapping out overfitted nonlinearities.

All three supervised methods were excellent at identifying important features using the optimal distance-based features (Fig. 2 D-E, Fig. S1 B). RF and KL often outperformed MLP, although all three supervised methods were close to perfect at identifying the important features for this setup. Note that to find good discriminators between states, one should use a classifier such as MLP or RF instead of KL divergence. However, seeing that a high KL divergence correlates with a proper choice of input coordinates suggests that it can be used as a *separation score* to describe how good a few selected features are at separating states.

For every method, we chose a set of hyperparameters to tune using cross-validation (**Tables S1-S6;** Fig. S4-S9) and included the results for the parameter set with best average performance in Fig. 2, S1 and S2. For PCA, we varied the number of components to consider in the aggregated importance per feature (**Table S1;** Fig. S4) and found that it is better to include all the PCs with corresponding eigenvalues (i.e. variance covered) within the same order of magnitude than to only include the first PC. For all three neural network based methods (RBM, AE and MLP) we evaluated different shapes of the hidden layers (**Table S2-S4;** Fig. S5-S7) and found that increasing the number of hidden nodes or layers does not necessarily increase performance in general. For the MLP and AE we considered the regularization parameter and for the RBM we evaluated the performance for different values of the learning rate. **For the AE we also varied the number of epochs used in training, the learning rate as well as the batch size.** Neither had a critical impact on the overall performance in this case, with the exception of having a too high learning rate equal to one for the RBM. We also computed the KL divergence with different discretization parameters (**Table S5;** Fig. S8) and trained RF classifiers as either a set of binary or multiclass classifiers with varying the number of estimators, the **maximal tree depth as well as the minimum number of samples to be a leaf** (**Table S6;** Fig. S9). A proper discretization size for KL seemed to be about 1% of the range of values covered in the data. **Tuning the number of minimum samples to be a leaf and** the number of estimators in RF classifiers tended to increase the accuracy to some extent. Otherwise, both of these methods were moderately sensitive to the choice of hyperparameters, although a set of binary RF classifiers were, unsurprisingly, considerably better at identifying important atoms of a specific state compared to a multiclass classifier, which was only able to select the important atoms for the entire ensemble. In general, we found that a small change in a method’s hyperparameter value for this application typically resulted in a small change in the importance profile, and that even a significantly different set of hyperparameter values performed with comparable accuracy.

**We note that it is unlikely that the best performing set of parameters in this study is optimal for all biological problems. In fact, it is unlikely that the optimal set of hyperparameters has been found even for the toy model.** Moreover, although important atoms in the toy model were successfully identified with distance based features, it is not sufficient to assume inter-residue distances to form an adequate set of input features for all biological problems. For example, it is possible that backbone dihedral angles are better indicators of certain types of conformational changes or that local root-meansquare deviation (RMSD) to experimentally resolved structures better describes how computationally sampled states agree with the experiments. To have a toolbox of methods which are not as sensitive to the choice of input coordinates in combination with metrics such as a separation score makes it straightforward to compare different sets of features and methods hyperparameters, and avoid making poor choices. Based on the learnings from the toy model benchmarks, we outlined our protocol in the form of a checklist (Box 1) that we followed as we transitioned into biological applications.

#### Box 1. Checklist for interpreting molecular simulations with machine learning.

1. Identify the problem to investigate
2. Decide if you should use supervised or unsupervised machine learning (or both)

a. The best choice depends on what data is available and the problem at hand
b. If you chose unsupervised learning, consider also clustering the simulation frames to label them and perform supervised learning
3. Select a set of features and scale them

a. For many processes, protein internal coordinates are adequate. To reduce the number of features, consider filtering distances with a cutoff
b. Consider other features that can be expressed as a function of internal coordinates that you suspect to be important for the process of interest (dihedral angles, cavity or pore hydration, ion or ligand binding, etc.)
4. Chose a set of ML methods to derive feature importance

a. To quickly get a clear importance profile with little noise, consider RF or KL for supervised learning. RF may perform better for noisy data.
b. For unsupervised learning, consider PCA, which is fast and relatively robust when conducted on internal coordinates
c. To find all important features, including those requiring nonlinear transformations of input features, also use neural network based approaches such as MLP. This may come at the cost of more peaks in the importance distribution
d. Decide if you seek the average importance across the entire dataset (all methods), the importance per state (KL, a set of binary RF classifiers or MLP), or the importance per single configuration (MLP, RBM, AE)
e. Chose a set of hyperparameters which gives as reasonable trade-off between performance and model prediction accuracy
5. Ensure that the selected methods and hyperparameter choice perform well under crossvalidation
6. Average the importance per feature over many iterations
7. Check that the distribution of importance has distinguishable peaks
8. To select low-dimensional, interpretable CVs for plotting and enhanced sampling, inspect the top-ranked features
9. For a holistic view, average the importance per residue or atom and visualize the projection on the 3d system
10. If necessary, iterate over steps 3-9 with different features, ML methods and hyperparameters

### Supervised and unsupervised learning pinpoint conserved residues and frequent target-protein binders of calmodulin

Calmodulin is a ubiquitous messenger protein activated by calcium ions (54). The Ca^2+^ -bound state typically resembles a dumbbell with the N-terminal and C-terminal domains that are separated by a fully helical linker (Fig. 3 A). When activated, clefts of hydrophobic residues are exposed in the two domains. The exposed hydrophobic surfaces enable binding to a large number of target proteins. Through regulation of these target proteins, CaM plays an important part in many physiologically vital pathways (55).

**Figure 3.**
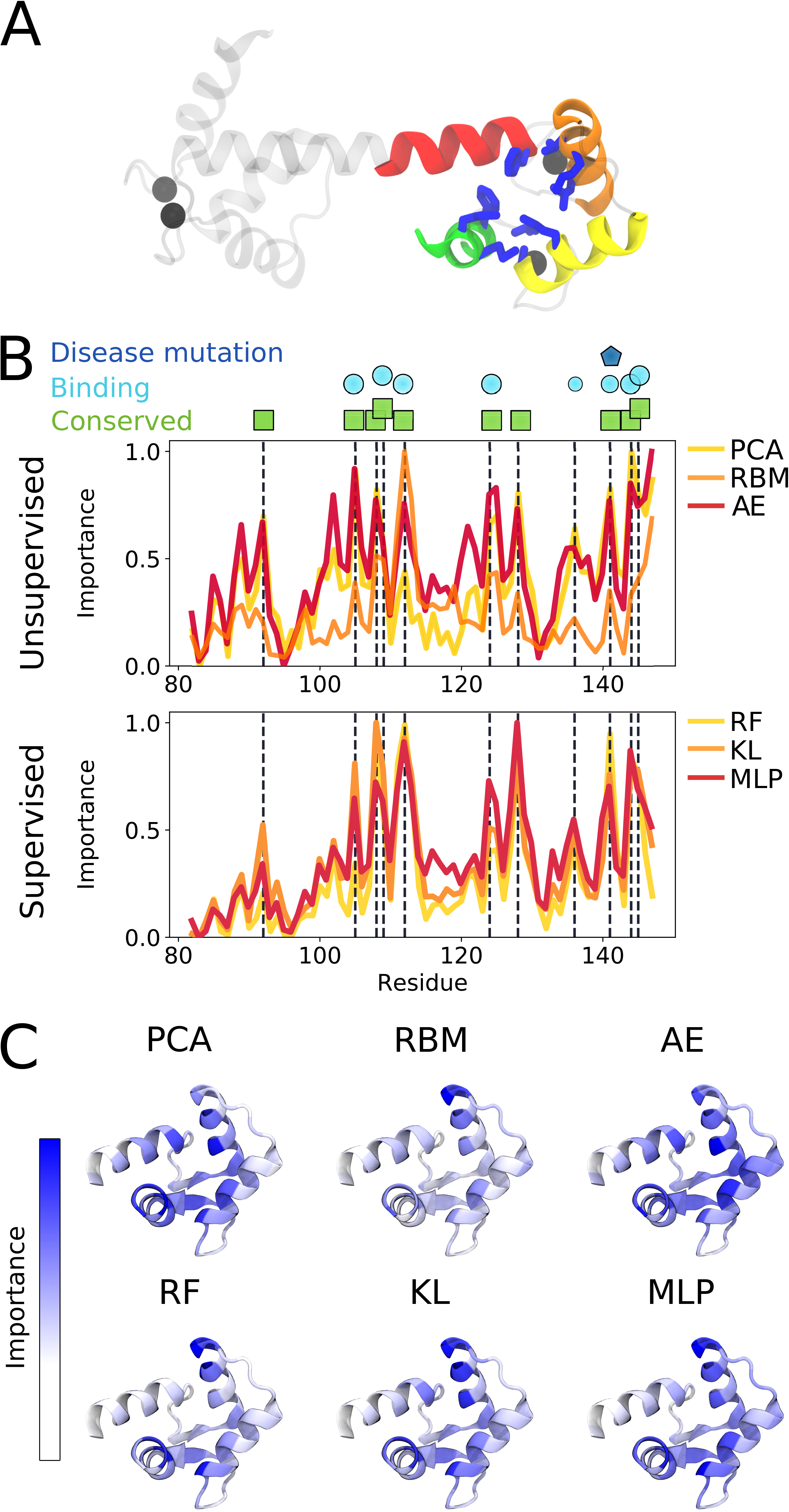
Important residues in the calmodulin C-terminal domain according to the feature extractors. (A) A molecular structure of Ca^2+^-bound CaM. The helices of the C-terminal domain are represented by colored ribbons, while Ca^2+^ ions are shown as black spheres. The residues identified as important in the analysis are shown with blue sticks. (B) The importance plotted along the residue sequence of unsupervised (upper) and supervised (lower) methods. The peaks identified through visual inspection of the supervised method results are marked by vertical dashed lines showing that most extractors agree. The peak residues are marked as being conserved (green box), involved in binding (cyan circle) or known to be mutated in a disease (blue pentagon). (C) Molecular structures of the CaM C-terminal domain with important residues identified by the six extractors highlighted in blue.

We made a qualitative comparison between different methods (extractors) applied to a dataset obtained with spectral clustering enhanced sampling trajectories of Ca^2+^-bound (holo) calmodulin (CaM). The dataset contains the six states of the C-terminal domain of CaM (47). **We hoped to use unsupervised and supervised methods to identify the biological descriptors underpinning the different states that were identified in a data-driven manner.**

Fig. 3 B and C show the average estimated feature importance obtained from the CaM dataset with six states, using internal coordinates as input. The peaks obtained from the different extractors are shown as sticks on the molecular structure (Fig. 3 A) and highlighted with vertical dashed lines in the importance profiles (Fig. 3 B). Notably, the supervised methods agree well while the unsupervised methods provide noisy profiles. Moreover, the residues that were found important are either highly conserved across phylogenetic groups (56), involved in binding target proteins (57), or known to cause severe cardiac diseases upon mutation (58). M109, M124 and M144-145, for example, are highly conserved (56) and recognized to play important roles for target-protein binding through various packings (59). Another example is F141, a well-studied mutation linked to long QT syndrome and decrease in Ca^2+^ affinity (58, 60). **We note that two peculiar peaks appeared in the AE profile of the CaM dataset; A102 and V121. The implications of these peaks and their dependence on AE architecture are discussed in Section S2.**

To monitor the dependency of the average importance profiles on the chosen number of states, a qualitative comparison between the RF importance profiles using five to eight states was carried out (Fig. S11). Evidently, the profiles obtained from six or more states overlap. In the five-state dataset, however, an extra peak appears at residue V121 (Fig. S11 B). This residue was specifically identified as important for state 2 and 4 in the five-state dataset, while only for state 6 in the six-state dataset (Fig. S12 A-C). Hence, it is assigned a large peak in the averaged profile of the five-cluster dataset and a smaller peak in the six-state dataset. We further note that both states 2 and 4 in the five-cluster dataset are poorly populated and merged into state 6 in the six-state dataset (Fig. S12 D-F). Therefore, the profiles with a smaller peak at V121, which was obtained with the six-or-more-state datasets appears more reasonable than that of the five-state dataset.

**Comparing the unsupervised and supervised profiles reveals two residues specifically identified by t**he supervised methods; V108 and A128. These two residues are not typical target protein binders. However, both residues are well conserved (56) and thus likely play stabilizing roles for exposing binding surfaces. In fact, the packing and repacking of V108 between F89 and F92 is linked to the transition between apo and holo-like C-terminal states (59). A third surprising peak is that of V136, which is not highly conserved (56) and only participates in binding to some extent (57). This residue is, however, part of the C-terminal domain beta-sheet that was shown to be disrupted in an extra-open state of the analyzed dataset (47). Thus, by contrasting the output of the different methods, it is evident that the supervised methods show more distinguishable peaks and the ensemble of methods successfully uncovers important residues of the CaM C-terminal domain.

### Supervised learning of the β2 adrenergic receptor identifies ligand interaction sites and reveals allosteric coupling, while unsupervised methods learn the main characteristics of GPCR activation

GPCRs form a large family of receptors in the human genome and act as an important class of modern drug targets (61). The β2 adrenergic receptor (β2AR; Fig. 4 A) is a class A GPCR often used as a prototype in studies of GPCR activation. In its native state without any binding partner, the intracellular part of the β2AR undergoes interconversion between an inactive and an active state allowing a G protein to bind to the intracellular region, a process called basal activity. Agonist ligands interact with residues in the orthosteric site and shift the equilibrium between states to make the active state more energetically favorable.

**Figure 4.**
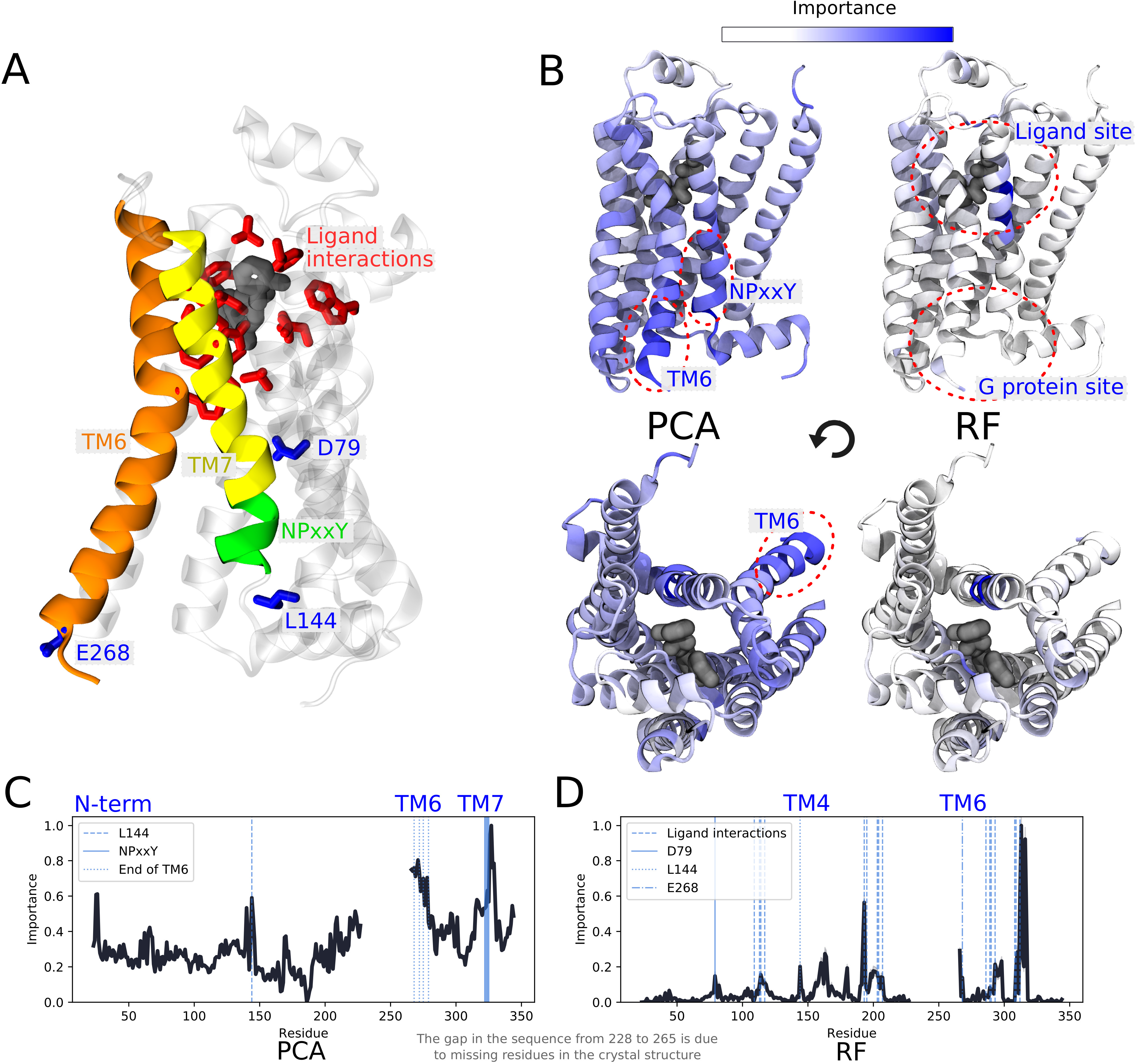
Important residues of the β2 adrenergic receptor using frames from a string-of-swarms simulation of the receptor’s activation (10). (A) A molecular structure of the β2 adrenergic receptor bound to an agonist ligand. Transmembrane helices TM6 and TM7 as well the NPxxY motif are represented as colored ribbons. Important residues known to interact with ligands are shown with red sticks while other key residues identified in the analysis are colored in blue. (B) active state structures with important residues highlighted in blue. (C) unsupervised learning with PCA. The G protein binding site undergoes the most significant conformational change upon activation, especially TM6 and the NPxxY motif at the bottom of TM7. (D) supervised learning with an RF classifier trained to identify protein conformations as either being ligand bound or ligand unbound. Residues known to interact with the ligand binding site (62) coincide with the residues identified as important. Important residues to discriminate between the two conditions are also located in the G protein binding site, an indication of allosteric coupling between the intracellular part of the receptor and the orthosteric site.

We analyzed the conformational ensemble of short trajectories from a string-of-swarms simulation of β2AR activation in the presence and absence of an agonist ligand (10). Unsupervised learning with PCA on the entire internal coordinate dataset revealed known hallmarks of activation (Fig. 4 B, C), including significant rearrangements of helices around the G protein binding site, especially the twist of the conserved NPxxY motif at the bottom of TM7 (63, 64). Another protruding hallmark of GPCR activation identified by unsupervised learning is the displacement of the intracellular part of TM6, including E268, which is the residue that undergoes the largest conformational change upon activation (65), as well as residues L272, L275 and M279 on TM6, whose side chains are rotated upon activation (66). The third most important region consisted of residues just below the extracellular N-terminus, a disordered region consistently changing its orientation throughout the simulation. Its large variance in interactions with other residues makes it prone to be identified as important by these methods, even though its movement is not considered a typical characteristic of GPCR activation. Finally, L144 at intracellular loop 2 was shown as important, and it is known to be implicated in β2AR/Gs coupling (67, 68).

Following the guidelines in our checklist (Box 1) we trained an RF classifier to discriminate between the ligand-bound and ligand-unbound simulation frames within the same dataset. After training, the most important residues were identified close to the ligand binding site (Fig. 4 B, D), most of which are well known to interact with agonist ligands (62). Other parts of the receptor interacting directly with these residues were also identified as important, especially residues along transmembrane helix 4 (TM4). T164, for example, interacts with ligand binding site residues S203, S207 and V114. In general, residues further away from the orthosteric site were considered less important (Fig. 4 D). One exception is the local peak at the D79 residue on TM2, the most conserved residue in all class A GPCRs. This residue is also thought to change protonation state during activation (Fig. 4 D) (10). Remarkably, important residues were further identified around the G protein binding site: L144 and E268 at the bottom of TM6, which sample both larger as well as smaller values of displacement in the apo trajectories (10). This result is indicative of allosteric coupling between the two sites and shows that the classifier is able to detect subtle local changes in the G protein binding site that are induced by agonist binding. It suggests that ligand binding not only shifts the energy of the active state, but forces the G protein binding site to adopt alternative conformations. The intracellular end of TM6 and the NPxxY motif do not appear as important on the other hand, presumably because they are observed in similar conformations both in the ligand bound and the ligand-unbound ensemble.

In Fig. 4 B-D it can clearly be seen that RF gives a more easily-interpretable profile compared to PCA, in which the peaks are distinct and the average importance is low. This is not merely a result of the chosen methods; it is also due to the kind of insights they provide. GPCR activation involves conformational change throughout the protein, while ligand binding and allosteric signalling are more local effects. To conclude, unsupervised ML was useful on this dataset to study the general large-scale activation mechanism, whereas supervised ML successfully showed the effect of agonist binding throughout the conformational ensemble.

### Supervised learning reveals state-dependent residues allosterically coupled to the gating charge

The voltage sensor domain (VSD) is a common module of voltage-gated ion channels responsible for reacting to changes in the membrane potential (69). It is usually composed of four transmembrane segments named S1 to S4, and a short surface C-terminal helix, the S4-S5 linker (Fig. 5 A) (70–72). A few positively charged residues on the S4 segment confer voltage sensitivity to the VSD function (Fig. 5 A). As they are located in a low dielectric medium, these residues are specifically sensitive to an externally applied electric field, and thus adapt their positions according to the field direction and strength (50, 73, 74). For instance, when the membrane is depolarized they move toward the extracellular solution. This motion engages the entire S4 segment which promotes the voltage sensor activation: the S4 transition from resting (inward-facing) to the activated (outward-facing) state (75–79). To ensure the stability of the positive charges inside the low dielectric medium, they are paired with negative charges of the three remaining segments. During activation, the positive charges switch negatively charged counterparts so that a network of salt bridges between S4 and S1-S3 is maintained in every conformational state of the VSD (75).

**Figure 5.**
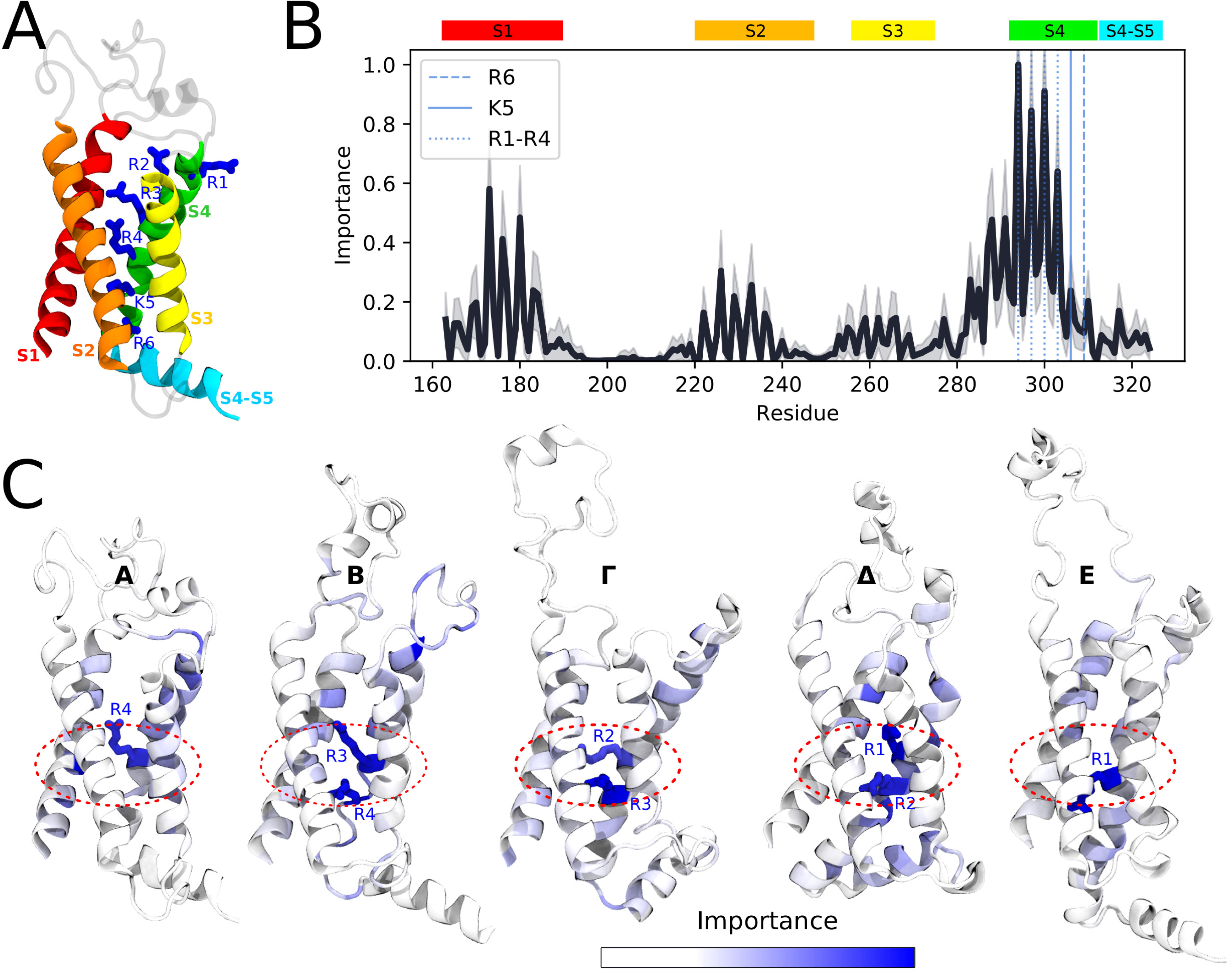
Identifying key residues for Kv1.2 voltage sensor function. (A) Cartoon representation of the Kv1.2 voltage sensor. The four membrane segments S1-S4 and the surface S4-S5 helix are shown. S4 positive charges are represented as blue sticks. (B) Per-residue importance obtained using RF classifiers and averaged over the five states of the Kv1.2 VSD. The colored rectangles denote the helical segments. S4 charges (R1, R2, R3, R4, K5 and R6) are marked with the blue dashed lines. The average importance and its standard deviation are shown as the black line and the shadow around it, respectively. (C) Per-state per-residue importance. The top-ranked residues are shown as sticks. The area of the hydrophobic plug is marked as a red dashed zone.

In a previous study, we reported five conformational states of the Kv1.2 VSD called E (resting), A (activated) and *Δ, Γ*, B (intermediate states) (49, 50). Here, we followed the provided checklist (Box 1) and trained a RF classifier to compare these states and elucidate their defining structural elements using more than 12,000 frames from an enhanced sampling trajectory (see Methods). The **simulation frames were extracted from basins in the free energy landscape** constructed by metadynamics. **Thus, even though the trajectory was not sampled from a Boltzmann distribution, the derived simulations snapshots represent conformations one would expect to see in an unbiased MD trajectory.** Fig. 5 B shows the per-residue importance obtained using RF classifiers, averaged over all five states. The first top-ranked residues (R1, R2, R3 and R4) are four out of six positive charges of the S4 segment (R1, R2, R3, R4, K5 and R6). Thus, in agreement with the classical view, the positively charged residues are suggested to be crucial in delineating the five conformational states of the Kv1.2 VSD. Analysis of the per-residue importance of each state (Fig. 5 C) reveals that while top-ranked charges are indeed state-specific (R4 in A; R3 and R4 in B; R2 and R3 in *Γ;* R1 and R2 in *Δ;* R1 in E), their localization is state-independent: all of them are placed near the so-called hydrophobic plug, the central part of the VSD where the response to the external electric field is the largest (50, 80, 81). Other highly ranked residues are located along transmembrane segments and facing the protein lumen. LRP performed on the MLP classifier further offers the possibility to attribute feature importance to each individual trajectory frame. Supplementary Movie 1 specifically shows per-frame feature importance profiles and provide comparable insight regarding the localization of important residues around the hydrophobic plug. Taken together, our data suggests that the interaction with the hydrophobic plug is a key aspect to distinguish between the metastable activation states in the Kv1.2 VSD.

### Supervised and unsupervised learning play different roles in identifying important features

Supervised and unsupervised methods are designed for different purposes and **the optimal choice of analysis method is case-dependent**. Sometimes the labels are inherent to the computational experiment, such as in the example of comparing ligand bound and ligand unbound simulations of β2AR, or when comparing the conformational ensemble of a wildtype protein with a specific mutant. For such cases, supervised learning techniques may reveal subtle differences invisible to unsupervised techniques. The performance evaluated with the toy model also shows that supervised learning generally outperforms unsupervised learning on labelled datasets.

Standalone unsupervised learning becomes more attractive when simulation data cannot be labeled directly. In order to use supervised learning for such cases, one would need to preprocess the data, for example by manual labelling or with a clustering method, to assign each frame to a state. Given adequate states, supervised learning will provide useful insights about the data, as was done for calmodulin and the Kv1.2 VSD in this study. **However, to arrive at biologically relevant insights requires the states themselves to contain useful information and it is important to bear this limitation in mind for practical applications.** Unsupervised learning techniques can also be used as prior feature extractors to supervised classifiers or provide a projection onto a low dimensional space where it is straightforward to cluster structures into states. A strength of having different tools for computing the importance per feature within the same framework is that it becomes straightforward to combine supervised and unsupervised learning to give a coherent view of what features are important in a system. It is simple to extend this framework to include more methods as long as there is an algorithm to derive the importance per input feature.

For practical applications, the best choice of method depends on the type of data available. Regardless, the results should be averaged over many runs and cross-validated to reduce the variance induced by stochastic solvers, as well as the risk of introducing inaccuracies due to overfitting. However, unless a good set of input features is known or can be validated, nonlinear ML methods such as neural networks will be more reliable than linear methods in general.

We also note that the protocol presented in this study is in itself not restricted to study data sampled from a Boltzmann distribution, as illustrated by the voltage-sensor domain application case. The biophysical conclusions one can draw from the analysis, on the other hand, will depend on the nature of the dataset. If one was to look for important features for an entire ensemble in a dataset containing highly biased samples, then taking the Boltzmann weight into account might aid interpretation. In general, this can be achieved by balancing the dataset according to the weight of the samples, and some models such as the Random Forest Classifier used in this study even support weighting every sample during training out-of-the-box.

### Feature selection and model parameter choice can enhance performance, but are not obstacles for interpretability

We used inverse interatomic distances as features for the biological systems, which puts more emphasis on local changes than mere distances, and normalized the values to enable the different ML methods to perform well (Methods). The choice of internal coordinates may seem straightforward following the results obtained from the toy model. For large systems, however, it may be computationally intractable to consider all internal coordinates as features and, in the case of limited amount of simulation frames, can also increase the chance of overfitting the feature extractors. In addition to performing crossvalidation to overcome this problem, distances which did not reach below a certain cutoff in the trajectory were pre-filtered in the case of CaM and VSD (Methods). This also has a physical significance since atoms close to each other have a stronger interaction energy. Long-range allosteric effects may still be discovered and propagated through nonlinear methods.

In principle, any intuitively interesting reaction coordinate can be included in the set of input features. In protein folding, dihedral angles are often reported as important, and should thus be included. Other collective variables such as hydration, ion binding, etc. may be considered as part of the input feature dataset. We note here too that we have only taken structural (and not dynamic) information into account, following the nature of the dataset we had at hand. To account for dynamics, for example a time-lagged autoencoder (11) could be used on continuous MD simulation trajectories, such that a neural network is trained to reconstruct features using features from previous times as input, and derive the resulting importance profiles from the trained model. In general, given a Markov State Model with a number of connected markov states (32, 33, 82, 83) our approach provides the means to make a data-driven interpretation on a molecular basis of the dynamics behind these states.

For supervised learning one should be aware of the difference between choosing a multiclass RF or MLP in contrast to using KL or RF with a one-vs-the-rest splitting of the data. One-vs-rest splitting in learning enables identifying important features for specific states as opposed to only reporting the important features across the entire ensemble (Fig. S7-S9). Furthermore, the difference between multiclass and one-vs-rest may have a major effect when dealing with unbalanced datasets. In this regard, algorithms such as LRP are powerful since they allow us to compute the feature importance for every individual frame (Supplementary Movie 1) and thereby enhance interpretability at a very fine-grained level.

Fig. S4-S9 demonstrate that the performance of the methods are affected by choosing different hyperparameters, such as the regularization constraints for a neural network, the size of the discretization parameters to compute the KL divergence **or the minimum number of samples to be a leaf in a RF classifier**. To include too many parameters in a model by adding an unnecessarily large number of hidden nodes in a MLP or to neglect parts of the dataset by considering too few PCs are both examples of what may reduce performance. We found it particularly challenging to obtain high accuracy with the AE and **it is possible** that additional effort in parameter tuning may improve its performance. Seeing that they are both unsupervised neural network based methods, it seems reasonable for the AE to obtain a performance comparable to the RBM, even though the AE used in this study, unlike the RBM or for example a variational autoencoder (84), does not make any strong assumptions about the distribution of input or hidden layer latent variables. Incidentally, an optimally trained AE with sigmoid activation and a single hidden layer is strongly related to PCA (85, 86). **For smaller sets of input features we do indeed observe a similar performance between PCA and the AE (Fig. S3). Thus, from a practical point view it is probably a more rewarding effort to reduce the number of input features, for example by using a distance-based cut off, than to expand the hyperparameter search space.** In general, neural network models have many tunable hyperparameters as well as a large number of weights that need to be fitted, while simpler methods such as PCA and KL have fewer parameters. This drawback of the complicated models needs to be taken into account for practical applications.

Putting some effort into crude parameter tuning should be a rewarding effort while excessive fine tuning may not reveal any new insights. Moreover, even though it is most likely possible to further improve the performance of these methods with extended efforts of parameter tuning, the optimal set of hyperparameters for a certain classifier likely depends on the choice of input features and the biological system. From a practical point of view, it seems instead to be more useful to follow the guidelines presented here using an adequate configuration of every ML method, ensure that it is robust to cross-validation and apply it to investigate the performance of different input features. Our checklist (Box 1), which covers the most important aspects of interpreting molecular simulations with ML (Methods), makes this process easier.

The fact that many models will yield the same conclusion irrespective of hyperparameter setup increases the credibility of these methods for practical purposes. For neural network based models, changing the network architecture leads to trained models with completely different underlying weight matrices. As shown in this study, networks of different shape and regularization constraints nevertheless identify the same important features in a simulation system. Although, similar arguments hold for other methods, this is a particularly important result due to the enormous interest and scepticism towards deep learning. This contradicts any criticism of the black-box nature of such methods: the biological properties propagated from the output to the input are clearly consistent and traceable, as well as interpretable.

## CONCLUSIONS

In this study we have applied various ML methods to identify important features from molecular simulations. We constructed a toy model that mimics real macromolecular behavior in order to perform a quantitative comparison between the methods and derive insights regarding their applicability for practical purposes.

When optimizing the performance of different ML methods, we found that the choice of proper input features had larger effect than hyperparameter tuning. Specifically, a great performance increase was observed when switching from raw Cartesian to internal distance-based features (potentially with a cutoff). Supervised learning methods, in particular, outperform unsupervised methods on a suboptimal Cartesian dataset. Furthermore, an MLP classifier was shown to be significantly better at identifying all important features of a system compared to RF or KL divergence. This indicates that a neural network is able to learn the transformation of input data to an optimal representation, and that more complex deep learning approaches are robust analysis tools when little, or no, prior knowledge is available to choose the input features. However, MLP’s output was arguably more difficult to interpret than that of the other supervised methods. For unsupervised learning on an optimal set of internal coordinates, all methods (PCA, RBM and AE) identified important residues with similar accuracies on smaller datasets. On larger datasets, however, PCA outperformed both RBMs and AEs with regards to accuracy as well as computational efficiency.

Finally, we successfully applied this protocol to derive key insights into three distinct types of biomolecular processes: the conformational rearrangements of the soluble protein calmodulin, the effect of ligand binding to a GPCR and the allosteric coupling of an ion channel voltage-sensor domain to a transmembrane potential. Rather than trying to identify one optimal method for all practical applications, we condensed our learnings into a protocol in the form of a checklist. In short, the ML methods were designed for different purposes and the best choice of analysis methods depends on the question at hand. Supervised learning should be used when applicable, for example for studying the small allosteric switches induced by ligand binding to a receptor or comparing distinct metastable states, whereas unsupervised methods can derive ensemble features of unlabelled data thus capturing the major conformational change of a protein.

**This work sheds new light on ML as a pillar in interpretation of biomolecular systems.** The possible applications of our approach spans from interpreting large amounts of simulation or experimental ensemble data to selecting CVs for enhanced sampling simulations. Compared to the computationally expensive MD simulations, these insights are cheap: all results in this study were computed on commodity hardware using software published in the public repository *demystifying* on github.

Humans did not evolve to interpret large sets of high dimensional data by mere visual inspection. ML methods were, on the contrary, specifically designed to process big datasets. Nevertheless, despite the increasing interest and faith put into ML, current state-of-the-art methods are not at a stage where computers are able to set up, run and analyze simulations autonomously. Instead we have shown how statistical models and algorithms commonly used to solve ML problems provide a powerful toolbox to efficiently make data-driven interpretations of biomolecular systems. **As the timescales accessible by simulations increase and the popularity of ML tools continue to thrive across many scientific areas, we anticipate that our approach can be useful to aid many researchers in demystifying complex simulations.**

## AUTHOR CONTRIBUTIONS

OF, MK and LD conceptualized the project. OF, MK and AW wrote the code and performed different parts of data curation and formal analysis: AW for CaM, OF for β2 and MK for the VSD. OF and MK designed and analyzed the toy model. LD supervised the project. OF, MK and AW wrote the first draft of the paper. All authors reviewed and edited the paper.

## ACKNOWLEDGEMENTS

This work was supported by grants from the Gustafsson Foundation and Science for Life Laboratory to LD. The simulations were performed on resources provided by the Swedish National Infrastructure for Computing (SNIC) at PDC Centre for High Performance Computing (PDC-HPC).

## Notes

https://github.com/delemottelab/demystifying

https://drive.google.com/open?id=19V1mXz7Yu0V_2JZQ8wtgt7aZusAKs2Bb

