## Supplementary text, methods and figures for "Molecular insights from conformational ensembles via machine learning"

**Running title: Molecular insights via machine learning**

**O. Fleetwood[1], M.A. Kasimova[1], A.M. Westerlund[1] and L. Delemotte[1]**

### SUPPORTING MATERIAL

[1] Department of Applied Physics, KTH Royal Institute of Technology, Science for Life Laboratory, Stockholm, Sweden

Supplementary Text

S1: Nonlinear displacement algorithm for constructing the toy model

Assuming that we have *M* states with *N* displaced atoms per state and that the magnitude of displacements is *d*, the states of the nonlinear toy model were defined as follows:

1. For every atom in state *m* = 0, …, *M*-1 and *n* = 0 ,.., *N-1* with position $\underline{r}_{n,m}$ :
    a. if n is 0: displace the atom with the vector {*md*, 0, *d(1-m)*}
    b. else:
        i. Rotate the atom around the z axis with the angle $(-1)^n d/r_{nn'}$ where $rr_{nn'} = |\underline{r}_{n,m} - \underline{r}_{n-1,m}|_{x,y}$
        ii. Rotate the atom around the z axis with the angle $(-1)^m d/r_{mm'}$ where $r_{nn'} = |\underline{r}_{n,m} - \underline{r}_{n-1,m}|_{y,z}$

### S2: Note on the extra peaks in AE of the CaM dataset and the effect on the importance profile depending on AE architecture

**Two extra peaks appeared in the AE profile of the CaM dataset; A102 and V121, Fig. 3 B. A102 is not highly conserved across phylogenetic groups, and although the mutation A102T is found in the Exome Sequencing Project human data, its phenotype is unknown (56). The two residues A102 and V121 interacted in states one to five, while in the sixth cluster, this interaction was disrupted (Fig. S13), and V121 was exposed to solvent due to a parallel organization of the C-terminal domain helices (47). In fact, A102 and V121 were specifically identified as important by RF for distinguishing state 6 from the others (Fig. S12 B, C). Due to the relatively small number of frames described by these features (Fig. S12 F), however, the residue importance is likely inflated in the AE profile.**

**To investigate whether these two peaks were due to an excessive AE compression, we compared the results obtained using two different network architectures; one layer with 100 hidden nodes as well as three with 300, 100 and 300 hidden nodes (Fig. S14). The obtained importance profiles are within standard deviations of each other which indicates that the compression using only one hidden layer is sufficient.**

Supplementary Tables

| Name | Values in this study |
| --- | --- |
| Number of components | Lowest value of the number of samples or the number of features (default) |
| Whiten | False (default; no whitening of the data was used) |
| SVD solver | ‘auto’ (default) |
| variance cutoff | 1 components, 2 components, 50 % of explained variance, 100% of explained variance or ‘auto’ (see Main text Methods) |

**Table S1. Hyperparameters for PCA. See the scikit-learn (41) documentation for a complete definition of these values.**

| **Name** | **Values in this study** |
|---|---|
| Number of hidden nodes | 1, 3, 10, 100, 200 |
| Learning rate | 0.001, 0.01, 0.1 (default), 1 |
| Batch size | 10 (default) |
| Number of iterations over the dataset | 10 (default) |

**Table S2. Hyperparameters for RBM.**

| Name | Values in this study |
|---|---|
| Hidden layer architecture | See Fig. S3, S6 and Main Text on Calmodulin for details |
| Activation function | Logistic sigmoid |
| Solver | ‘adam’ (default) |
| Alpha (regularization term) | 1e-2, 1e-3, 1e-4 (default), 1e-5 |
| Batch size | ‘auto’ (default; same as min(200, number of samples)), 10, 100 |
| learning_rate_init | 0.001 (default) |
| Maximum number of iterations/Number of epochs | 20, 200 (default) |
| Shuffle samples in each iteration | True |
| Tolerance | 1e-2, 1e-4 (default) |
| Early stopping | True (default) |
| Beta1 (parameter specific for the adam solver) | 0.9 (default) |
| Beta_2 (parameter specific for the adam solver) | 0.999 (default) |
| Epsilon (parameter for numerical stability in the adam solver) | 1e-8 (default) |
| Maximum number of epochs to not meet tolerance improvement | 10 (default) |

**Table S3. Hyperparameters for AE.**

| Name | Values in this study |
|---|---|
| Hidden layer architecture | [100] (default), [1000], [10], [50x10], [30x10x5] |
| Activation function | Relu (default) |
| Alpha | 1e-1, 1e-2, 1e-3, 1e-4 (default) |
| Batch size | ‘auto’ (default) |
| Maximum number of iterations/Number of epochs | 100000, 200 (default) |
| Tolerance | 1e-4 (default) |
| One versus the rest classification | False |

**Table S4. Hyperparameters for MLP. For additional network parameters, the same values as in Table S3 were used.**

| Name | Values in this study |
| --- | --- |
| bin width | 0.01, 0.1, 0.2, 0.5 and ‘auto’ (taken as the standard deviation of the dataset) |
| One versus the rest classification | True, False |

**Table S5. Hyperparameters for KL.**

| Name | Values in this study |
|---|---|
| Number of trees in the forest | 10, 100 (default), 200, 1000 |
| Method to measure the quality of a split | 'gini' (default) |
| Max depth of a tree | No limit (default; expand trees until the nodes are pure or to the minimum samples required to split a node), 1, 10, 100 |
| Minimum samples required to split a node | 2 (default) |
| Minimum samples required to be at a leaf | 1 (default), 0.25 times the number of samples, 0.45 times the number of samples |
| Number of features considered looking for the best split | 'auto' (default; the square root of the number of input features) |
| Bootstrap samples to build trees | True (default) |
| Class weight | Not provided, meaning that all classes had equal weights when building the trees |
| One versus the rest classification | True, False |

**Table S6. Hyperparameters for RF.**

Supplementary Figures

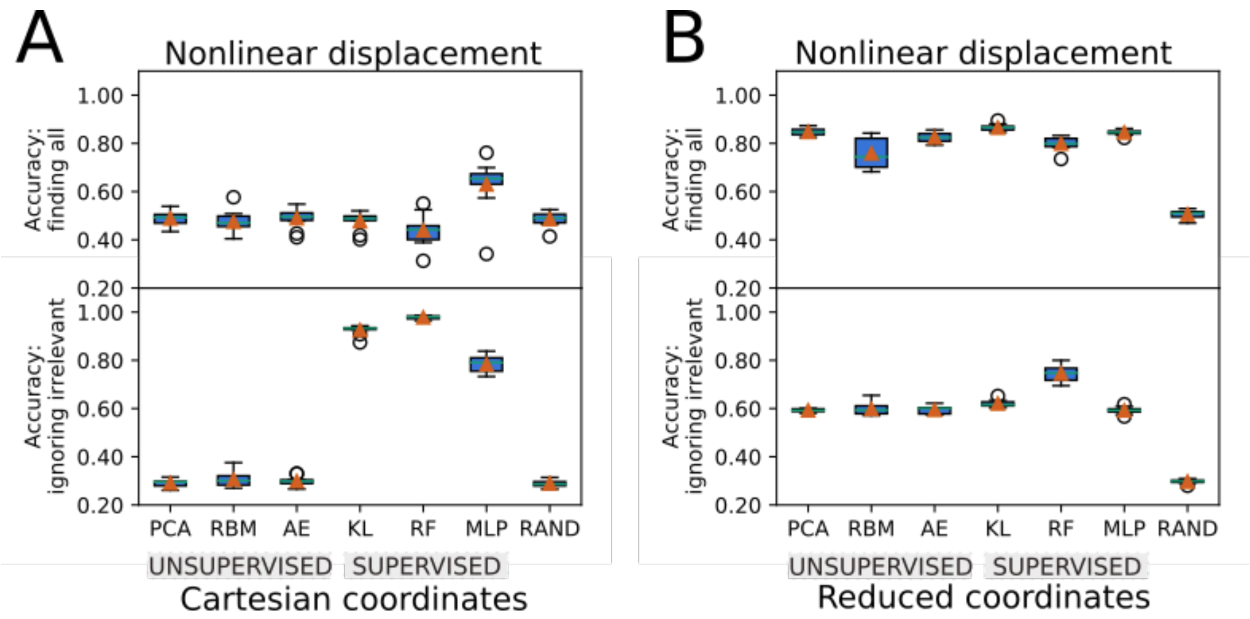


**Figure S1. Nonlinear displacement toy model used to benchmark the ML methods' ability to extract important features.**

**(A), (B) Boxplots showing the accuracy of the different methods using either flattened Cartesian coordinates (A) or a reduced set of inverse interatomic distances as features (B). The best performing set of hyperparameters found after benchmarking (Fig. S4-S9) have been used.**

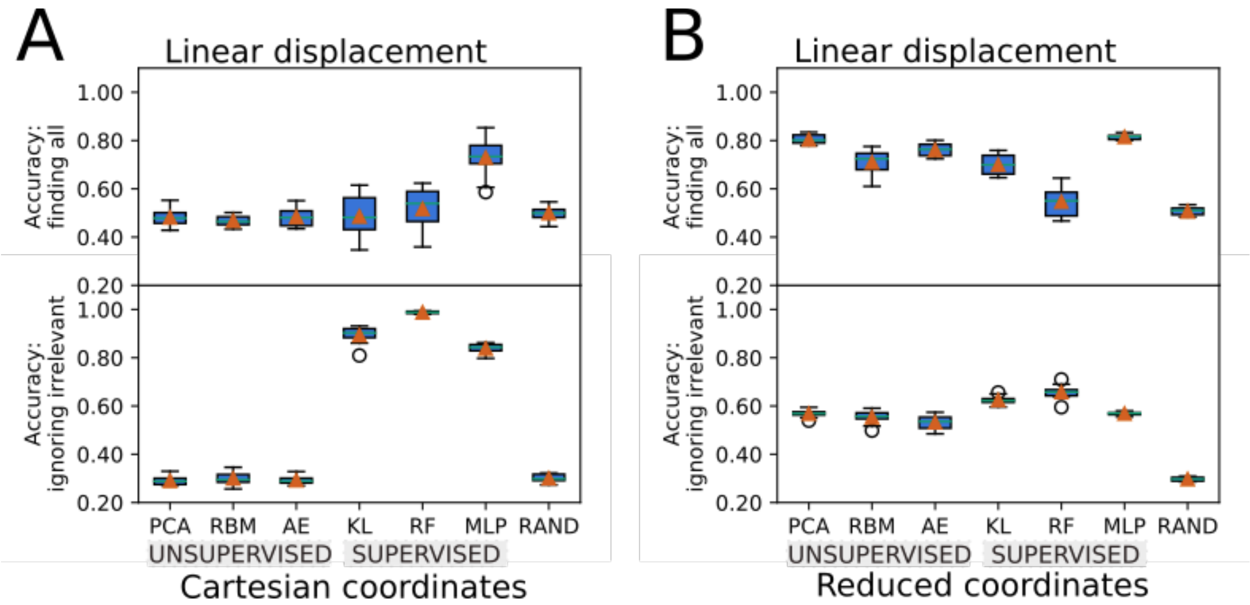


**Figure S2. Toy model used to benchmark the ML methods' ability to extract important features, using a noise level equal to 50 percent of the strength of the displacement.**

**(A), (B) Boxplots showing the accuracy of the different methods using either flattened Cartesian coordinates (A) or a reduced set of inverse interatomic distances as features (B). The best performing set of hyperparameters found after benchmarking every method (Fig. S4-S9) have been used.**

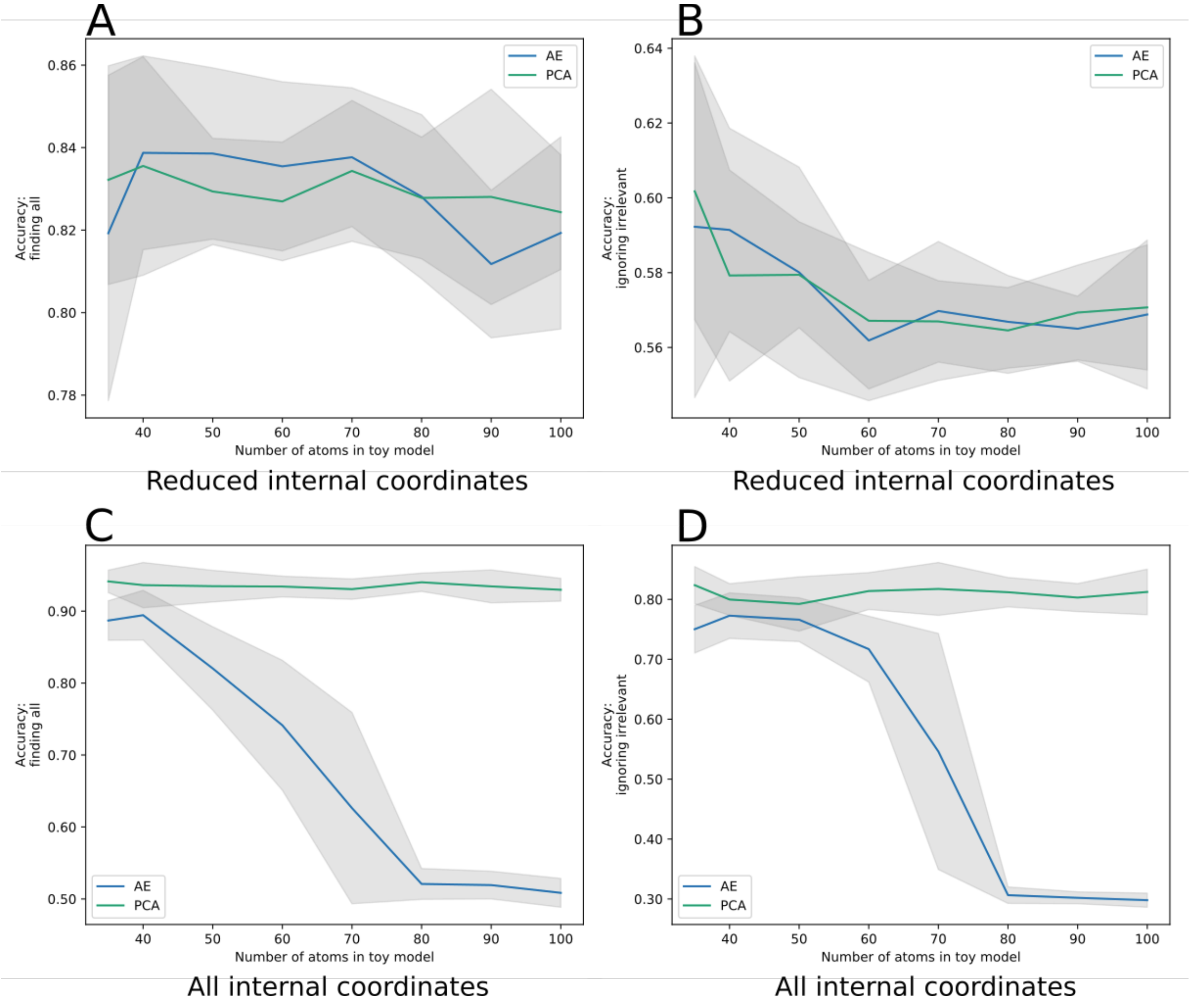


**Figure S3. Comparison of the accuracy of PCA and AE applied to the toy model for varying number of number of atoms in the system. (A), (B) for a reduced set of internal coordinates. (C), (D) for the full set of internal coordinates. The shaded area spans the standard deviation taken from 10 instantiations of the toy model. In all simulations, the toy model contained three clusters defined by the linear displacement of 10 % of the atoms in the system. The compression and decompression layers in the AE contained 50 % and 25 % of the input features. For systems greater than 40-50 atoms, the AE was significantly outperformed by PCA on the full set of internal coordinates.**

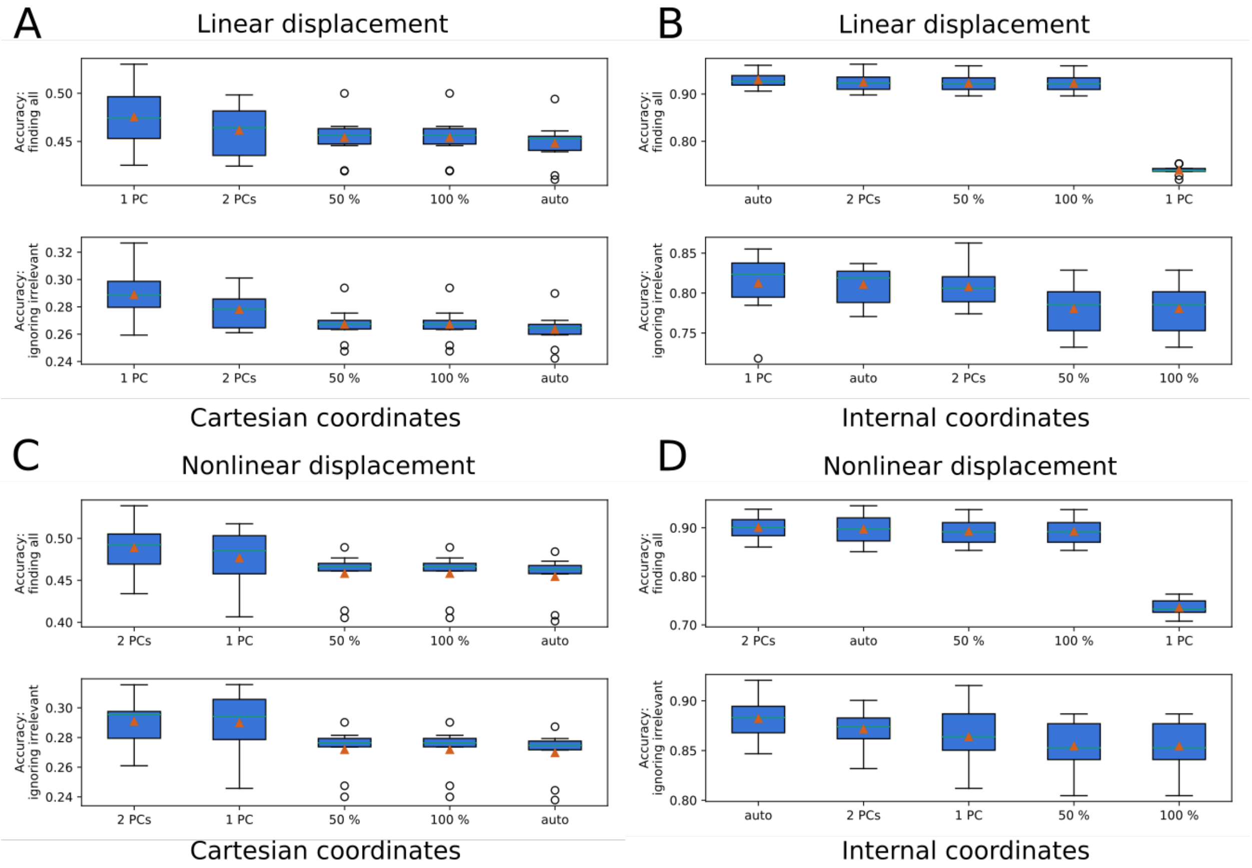


**Figure S4. Hyperparameter tuning of PCA applied on the toy model.**

**The number of PCs to include has either been determined by automatically finding ('auto') the first PC for which the next corresponding eigenvalue differs by at least an order of magnitude, by selecting a fixed number of PCs or by summing the variance covered by PCs until it exceeds a threshold given in percent. (A) Using cartesian coordinates and linear displacements, (B) using inverse distances and linear displacements, (C) using cartesian coordinates and nonlinear displacements, and (D) using inverse distances and nonlinear displacements.**

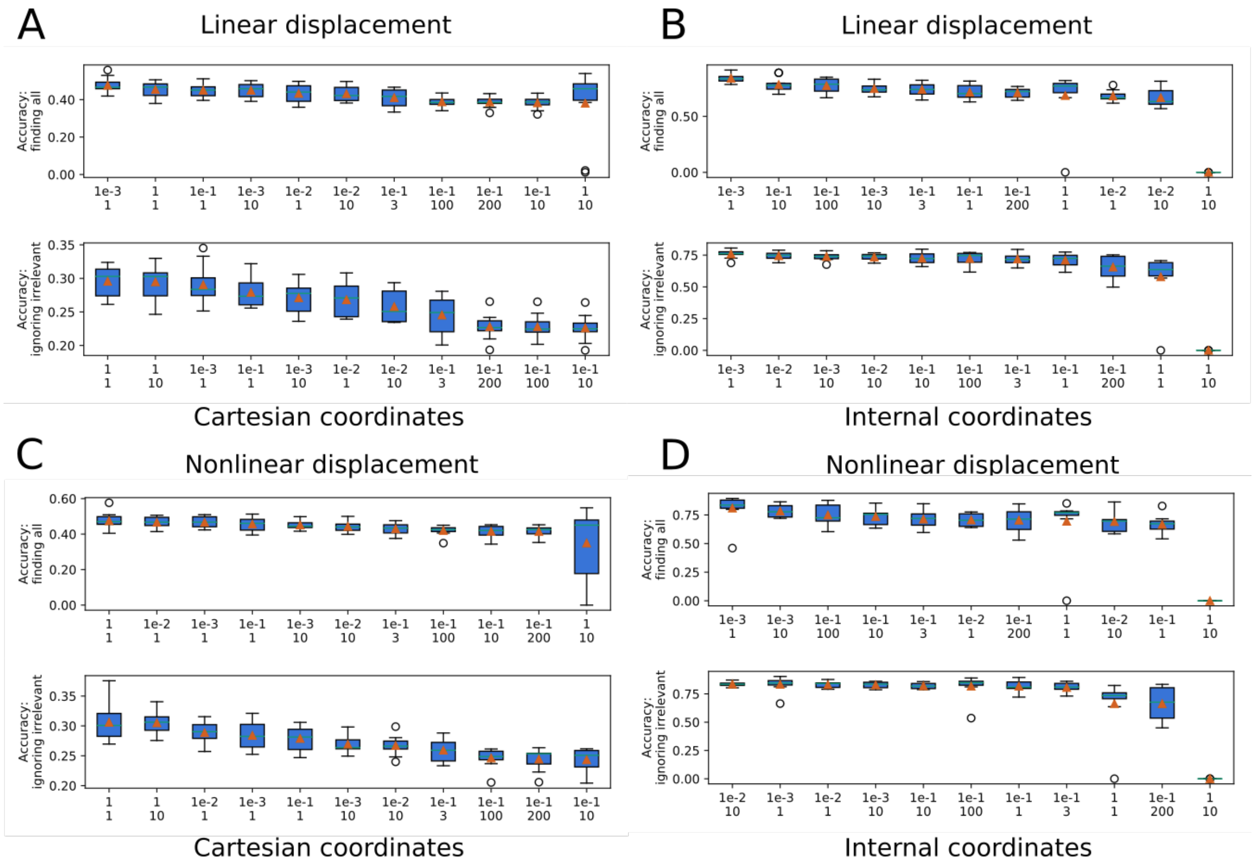


**Figure S5. Hyperparameter tuning of RBM applied on the toy model for different learning rates (first row on x-axis) and number of nodes in the hidden layer (second row on x-axis).**

**(A) Using cartesian coordinates and linear displacements, (B) using inverse distances and linear displacements, (C) using cartesian coordinates and nonlinear displacements, and (D) using inverse distances and nonlinear displacements.**

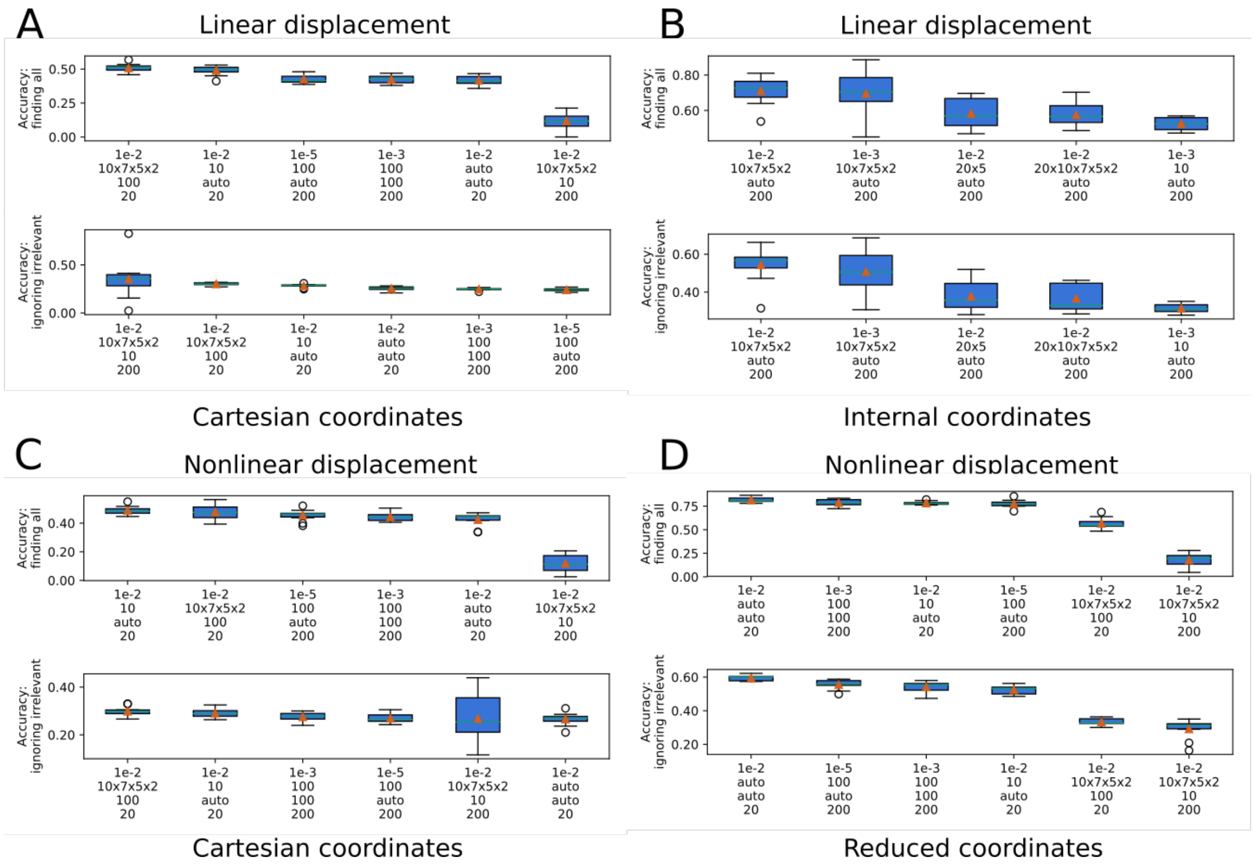


**Figure S6. Hyperparameter tuning of AE applied on the toy model for different combinations of regularization parameters (first row on x-axis), hidden layer architectures (second row on x-axis; auto corresponds to layers of 50% and 25% of the size of the input features), the batch size (third row) and maximum iterations (fourth row; also referred to as the number of epochs).**

**(A) using cartesian coordinates and linear displacements. (B) using inverse distances and linear displacements, only with the default values of the batch size and the maximum number of iterations. (C) using cartesian coordinates and nonlinear displacements. (D) using a reduced set of inverse distances and nonlinear displacements.**

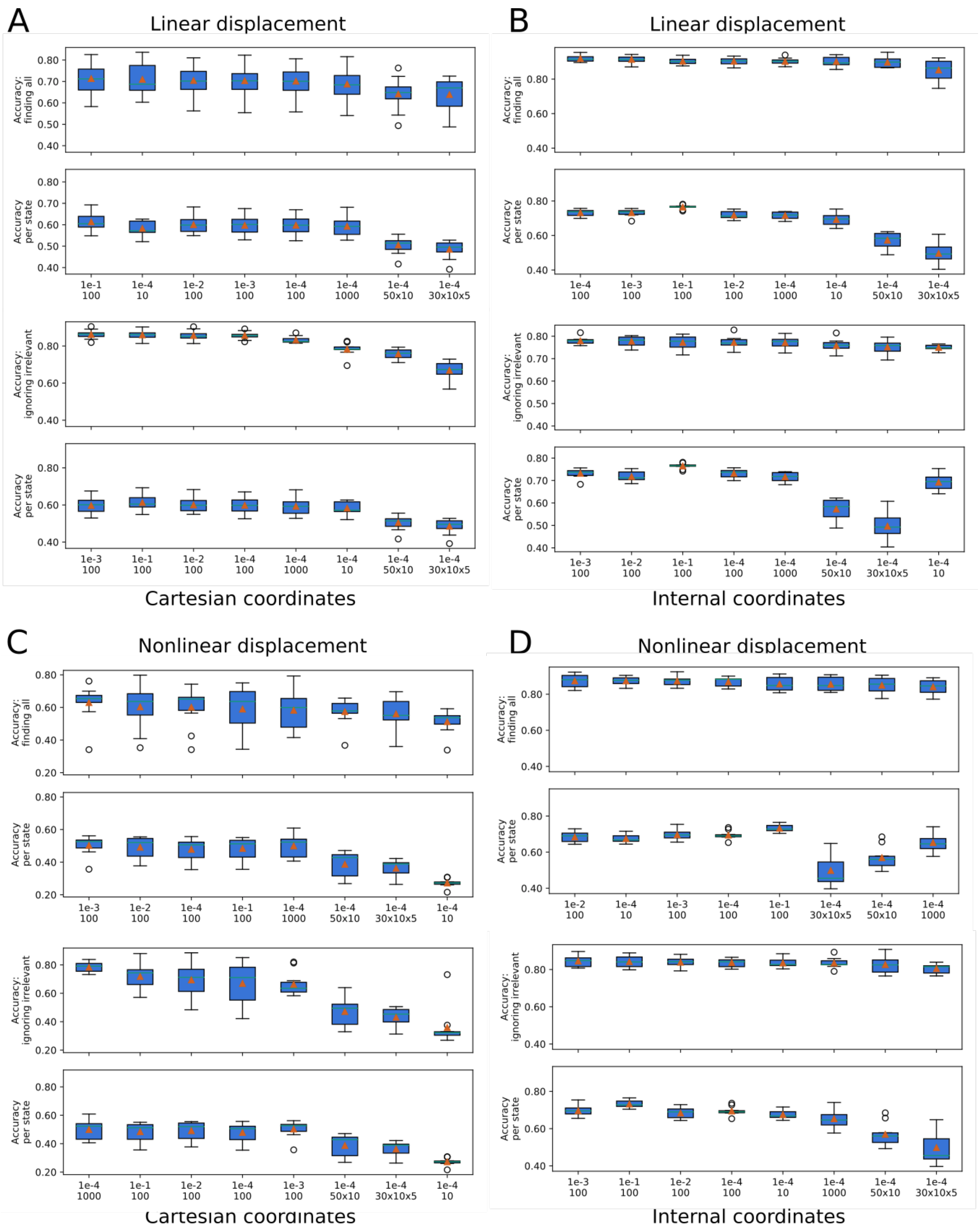


**Figure S7. Hyperparameter tuning of MLP applied on the toy model for different combinations of regularization parameters, α, (first row on x-axis) and hidden layer architectures (second row on x-axis).**

**(A) Using cartesian coordinates and linear displacements, (B) using inverse distances and linear displacements, (C) using cartesian coordinates and nonlinear displacements, and (D) using inverse distances and nonlinear displacements. The accuracy at identifying all displaced atoms in all states as well as the accuracy at identifying displaced atoms in a specific state is shown.**

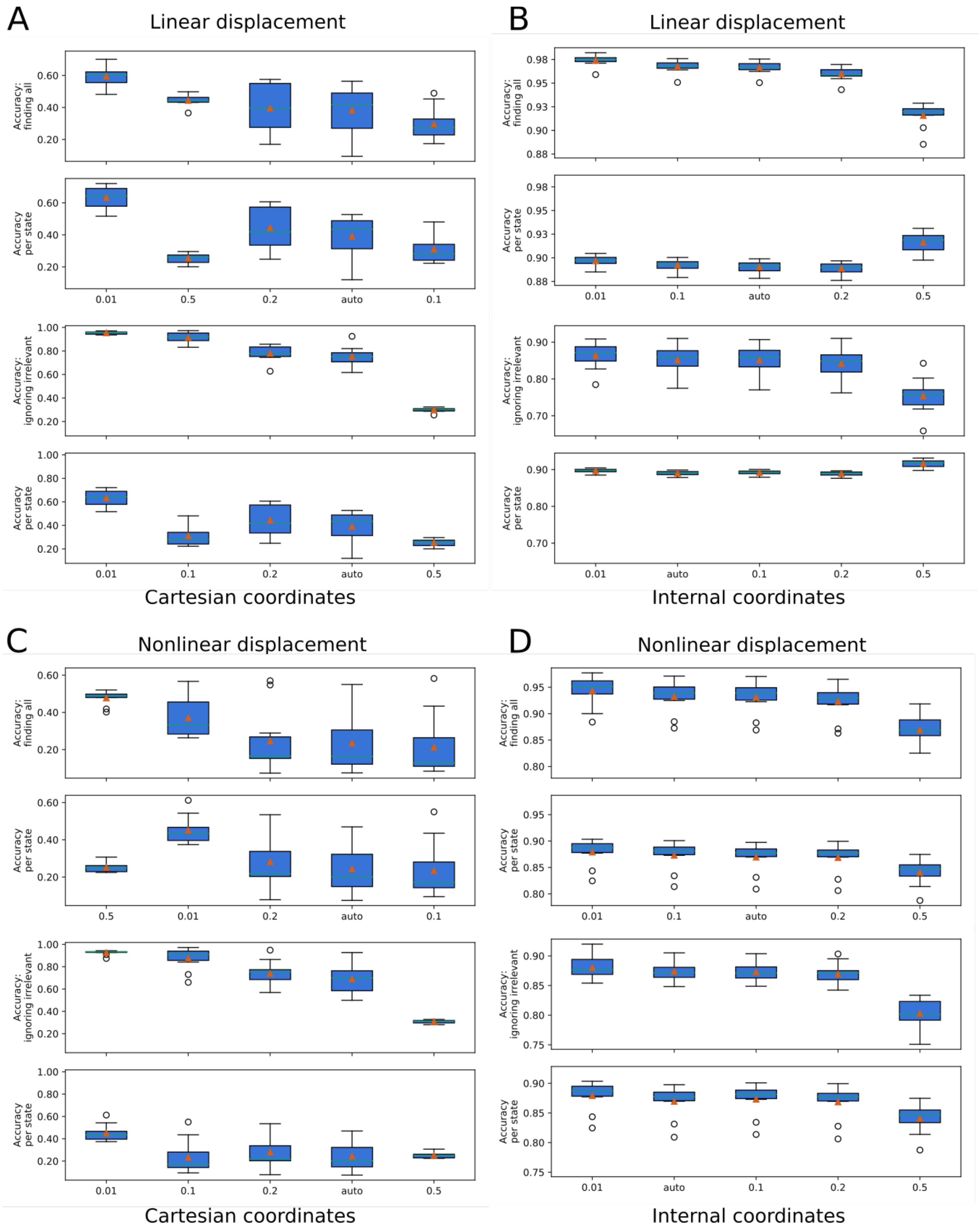


**Figure S8. Hyperparameter tuning of KL divergence applied on the toy model for different bin sizes used for discretization.**

**(A) Using cartesian coordinates and linear displacements, (B) using inverse distances and linear displacements, (C) using**

**cartesian coordinates and nonlinear displacements and (D) using inverse distances and nonlinear displacements. The accuracy at identifying all displaced atoms in all states as well as the accuracy at identifying displaced atoms in a specific state is shown.**

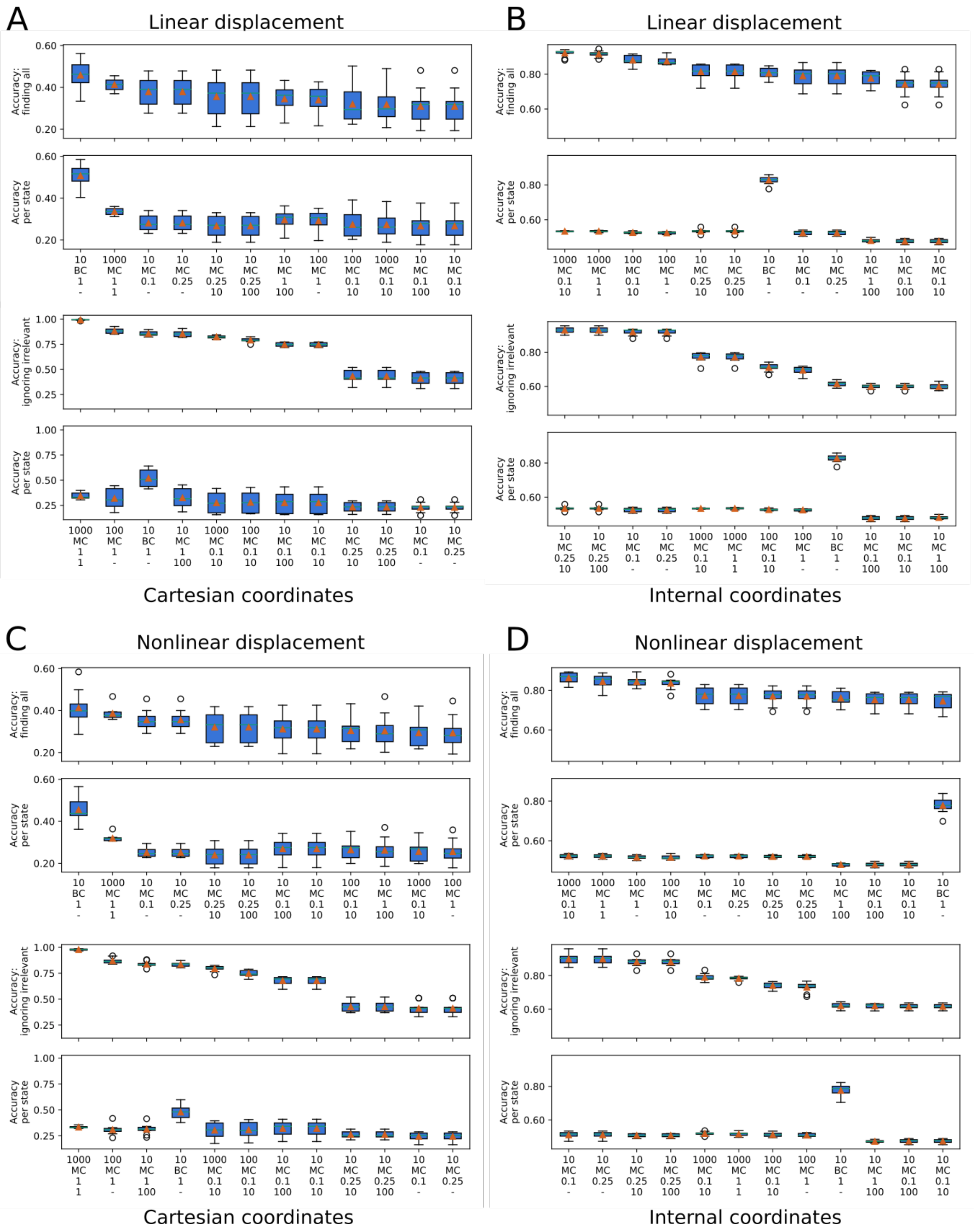


**Figure S9. Hyperparameter tuning of RF applied on the toy model for different number of estimators in the model (first row on x-axis), if it was trained as a single multiclass classifier (MC) or a set of binary classifiers (BC; second row on x-axis), for**

**different number of minimum samples required to be a leaf (third row; fractional number should be multiplied with the number of samples to obtain an integer limit) and for different values of the max depth (fourth row; if undefined the trees are expanded until all leaves are pure).**

**(A) Using cartesian coordinates and linear displacements, (B) using inverse distances and linear displacements, (C) using cartesian coordinates and nonlinear displacements, and (D) using inverse distances and nonlinear displacements. The accuracy at identifying all displaced atoms in all states as well as the accuracy at identifying displaced atoms in a specific state is shown.**

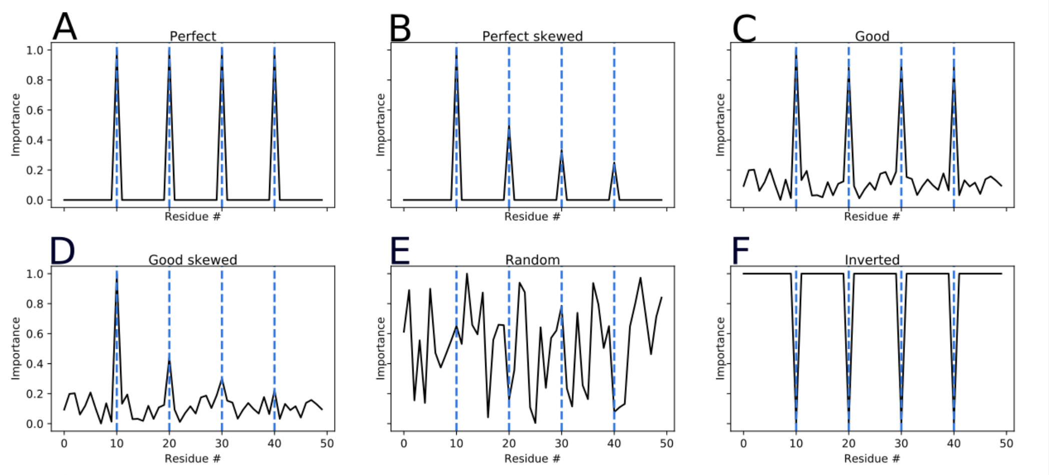


G

| Type | Name | Finding all | Ignoring irrelevant |
|---|---|---|---|
| A | Perfect | 1.00 | 1.00 |
| B | Perfect skewed | 0.77 | 1.00 |
| C | Good | 0.91 | 0.43 |
| D | Good skewed | 0.66 | 0.29 |
| E | Random | 0.15 | 0.06 |
| F | Inverted | 0.00 | 0.00 |

**Figure S10. Accuracy score values for example distributions.**

**The vertical dashed lines highlight the residues of high importance in the true distribution. (A) A distribution of perfect accuracy. All relevant residues have an importance equal to 1 and all other residues have 0 importance. (B) A skewed version of the perfect distribution. All irrelevant residues have 0 importance and the relevant residues have varying peak heights. (C), (D) The perfect distributions in (A) and (B) with random noise added to the distributions. (E) A distribution of random noise between 0 and 1. (F) The inverse of the true distribution has zero accuracy for both metrics. (G) Summary of the scores for all distributions. The mse-based method 'finding all' is sensitive to peak height whereas the 'ignoring irrelevant' accuracy method is not. On the other hand, the 'ignoring irrelevant' method is more sensitive to noise.**

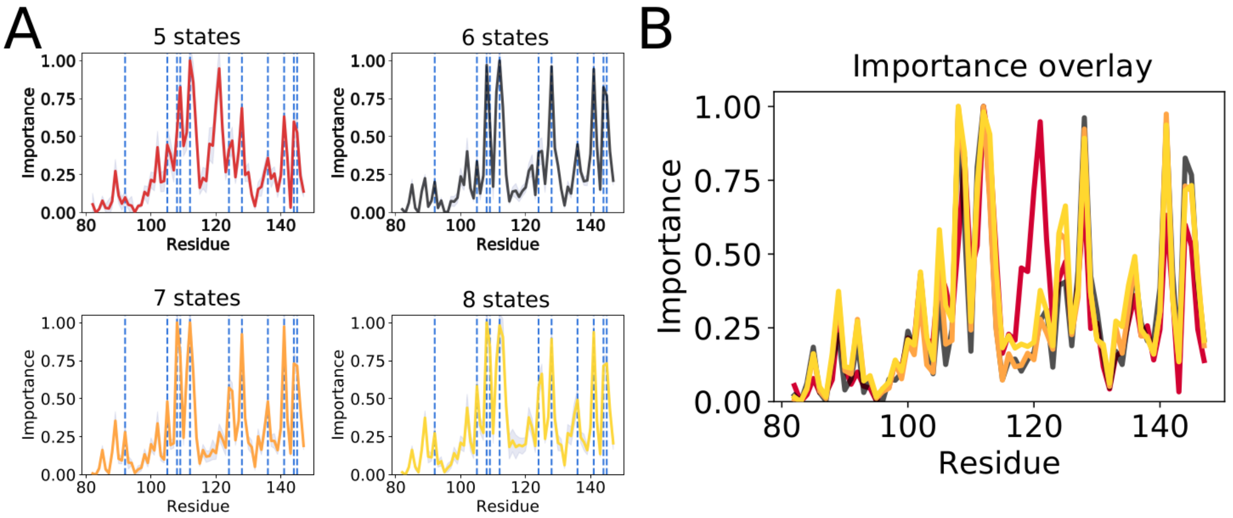


**Figure S11. Importance profile of RF applied to CaM datasets of varying number of clusters.**

**(A) The individual importance profiles with standard deviation. The peaks that are common across estimators are marked by blue vertical dashed lines. (B) The average importance profiles overlay to highlight similarities between the obtained profiles.**

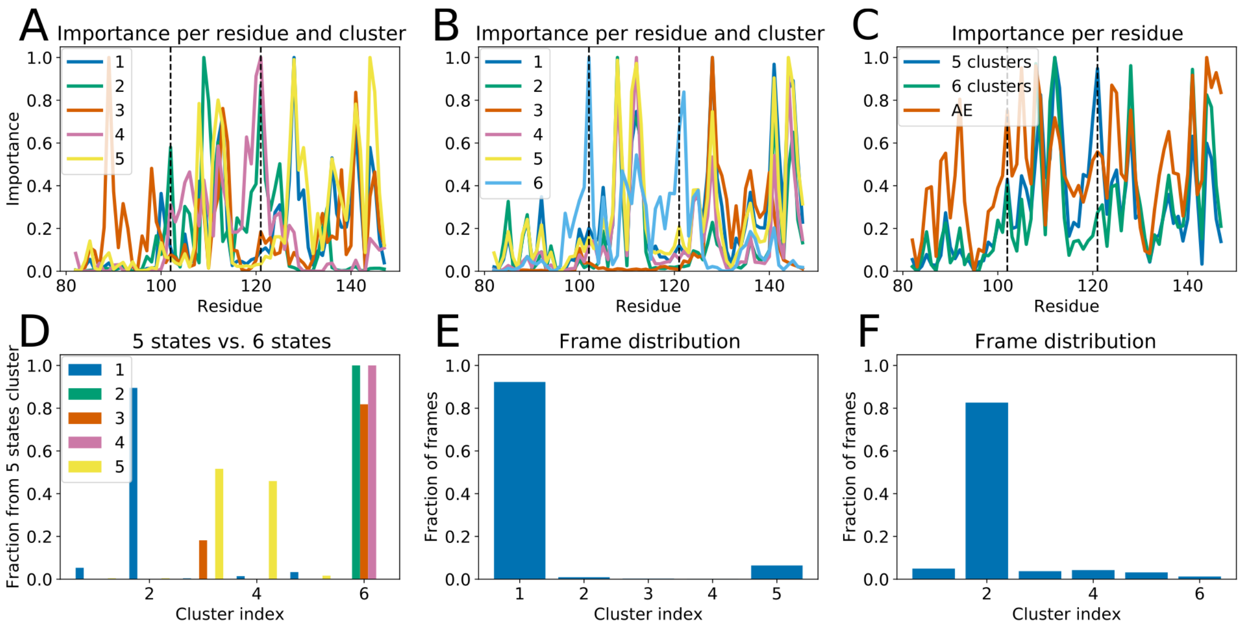


**Figure S12. Comparing the importance profile of RF applied to CaM datasets of five and six states, as well as the actual clustering.**

**(A) The importance per residue and cluster of the five-cluster dataset. (B) The importance per residue and cluster of the six-cluster dataset. (C) Average importance per residue of AE, five- and six-cluster datasets overlayed to highlight similarities between the obtained profiles. Residues A102 and V121 are marked by black vertical dashed lines. (D) The split of five-state dataset states into the six-state dataset. The bars correspond to the fraction of a five-cluster state that was clustered six-state dataset. Each color represents a state in the five-state dataset. (E) The fraction of frames in each state of the five-cluster dataset. (F) The fraction of frames in each state of the six-cluster dataset.**

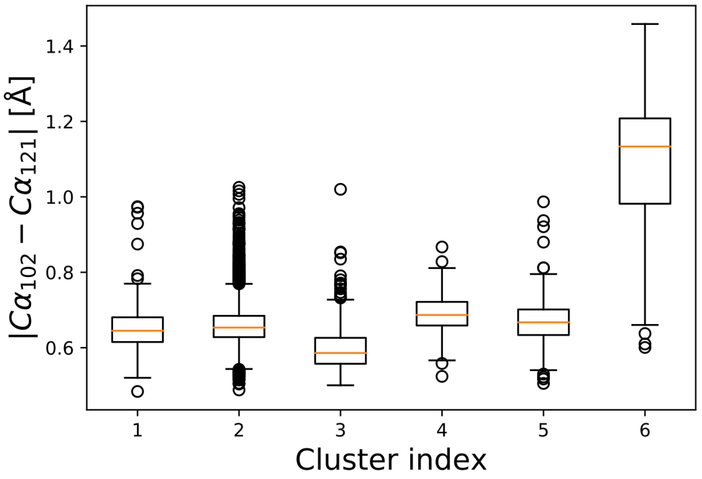


**Figure S13. Distribution of distances between A102 and V121 C-alpha atoms in the six CaM states identified with spectral clustering.**

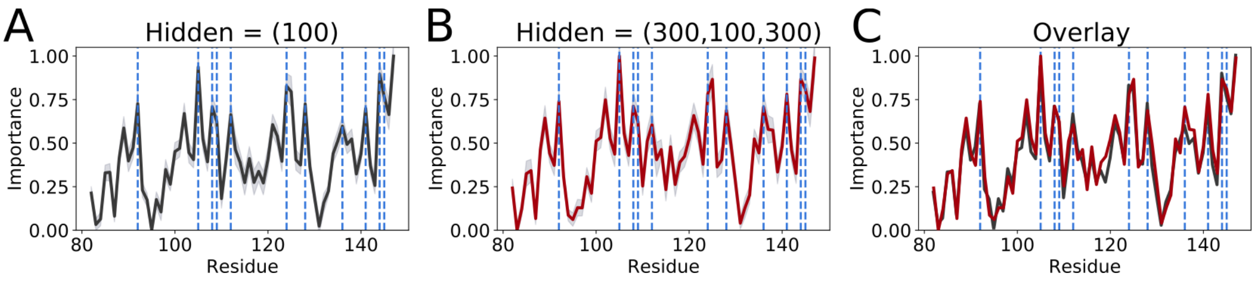


**Figure S14. Importance profile of AE applied to the CaM dataset, with varying the number of hidden nodes.**

**The individual importance profile with standard deviation using (A) one hidden layer of size 100, and (B) three hidden layers of size 300, 100 and 300. The peaks that are common across estimators are marked by blue vertical dashed lines. (C) The average importance profiles overlay to highlight similarities between the obtained profiles.**
